# Reprogrammed SimCells for Antimicrobial Therapy

**DOI:** 10.1101/2025.11.15.688600

**Authors:** Yun Dong, Xianglin Ji, Tao Dong, Yun Wang, Erik Bakkeren, Kevin R. Foster, Wei E. Huang

## Abstract

Antimicrobial resistance (AMR) is a critical global health challenge. In this study, we developed a novel platform based on chromosome-free and non-replicating simple cells (SimCells, size 1-2 µm) and mini-SimCells (size 100-400 nm) for targeted pathogen elimination. Engineered with surface-displayed nanobodies, SimCells and mini-SimCells selectively bind bacteria expressing specific antigens (e.g., OmpA in *E. coli*). The selective interactions facilitate close SimCell-pathogen proximity, enabling two antimicrobial mechanisms: direct injection of toxic effectors into bacterial cytoplasm via a heterologous expression of type VI secretion system (T6SS), and enzymatic conversion of aspirin into catechol by engineered salicylate hydroxylase, leading to sustained local production of hydrogen peroxide (H₂O₂). Our results demonstrate that both reprogrammed SimCells and mini-SimCells can eliminate target *E. coli* with high specificity and efficiency. Multi-dose reprogrammed mini-SimCell treatment led to a 10³-fold selective reduction of targeted bacteria in mixed microbial communities, with minimal disruption to non-target bacteria. We demonstrate that reprogrammed mini-SimCells, engineered with nanobody targeting outer membrane protein OmpA of the clinically relevant multidrug resistant pathogen *E. coli* ST131, achieved elimination efficiencies over 97% at 24 and 48 hrs. This modularised ‘plug-and-play’ antimicrobial platform provides a highly specific, efficient and adaptable solution for combating diverse AMR pathogens.

**Significance Statement:** Antimicrobial-resistant (AMR) bacteria cause millions of deaths worldwide annually, representing a critical global health challenge. In this study, we developed a novel therapeutic platform using two types of chromosome-free, non-replicating engineered bacterial cells: SimCells (1–2 µm) and mini-SimCells (100–400 nm). These SimCells were engineered to selectively bind to targeted *E. coli*, facilitating precise delivery of toxic proteins via a Type VI secretion system (T6SS) and localised generation of hydrogen peroxide from aspirin. We demonstrate that mini-SimCells eliminated over 97% of a targeted AMR strain within 48 hours. Moreover, multiple-dose administration achieved a selective 10³-fold reduction of targeted *E. coli* in mixed microbial communities. This modular ‘plug-and-play’ platform offers an adaptable solution against diverse multi-drug-resistant pathogens.

## Introduction

Antimicrobial resistance (AMR) is one of the top global public health challenges facing humanity by the World Health Organization (WHO)(1, 2). Predicted to become the next worldwide pandemic, AMR could cause over 10 million deaths annually and 100 trillion dollars of global GDP loss cumulatively by 2050(3–6). Despite this urgency, the discovery of new antibiotics has stagnated. The last new class of antibiotics was discovered in the 1980s(7, 8), with only recent reports in 2024 announcing a promising new antibiotic, zosurabalpin(9). Significant advancements in chemical synthesis(10, 11), high-throughput screening(12–14), rational design(15–17), and particularly artificial intelligence (AI)(18–23), have provided new pipeline for new antibiotic discovery. However, the antibiotic discovery might be insufficient compared to the rapid pace and spread of AMR. It is alarming that AMR has been developed for every approved antibiotic class currently being launched(24). The development of alternative antimicrobial strategies has been promoted, including therapies based on bacteriophage(25–27), monoclonal antibodies(28–30) and antibacterial peptides(31–33). However, these approaches face limitations such as stability issues, potential toxicity, and high manufacturing costs(33–35) (Supplementary Table S1).

Advancements in synthetic biology have led to the design of genetically engineered live microorganisms for the treatment or prevention of AMR pathogens in humans(36). Various microbial genetic components and tools have promoted the development of live biotherapeutic products (LBPs), providing a promising approach to address metabolic disorders, cancer therapy, and other unmet clinical needs(37, 38). However, biocontainment and biosafety concerns still restrain bacterial therapeutics from reaching a high public acceptability(39). To mitigate safety concerns, probiotic strains such as *Lactococcus lactis*(40) and *Escherichia coli* Nissle 1917 (EcN)(41, 42) have been used as chassis for therapy. Alternatively, attenuated strains with native tropism or colonising features, such as *Listeria*(43) and *Salmonella*(44, 45), are also employed. Despite the selection of the safety profile of the chassis, the risk of uncontrolled replication remains a challenge. A few strategies, such as conditional kill switches(46) and auxotrophy(47), have been used to prevent engineered bacteria from proliferating uncontrollably within or outside the human body and from escaping into the environment. Although these measurements restrict growth, concerns persist about the evolutionary stability of the introduced genetic modification and the potential risk of horizontal gene transfer (HGT).

To address these challenges associated with bacterial AMR, we developed customisable platforms based on SimCells (simple cell)(48) and mini-SimCells(49, 50) for use in antimicrobial therapy. SimCells and mini-SimCells are chromosome-free, non-replicating and reprogrammable bacterial chassis that function as “smart” bioparticles. SimCells (size 1-2 µm) are produced by removing native chromosomes using tightly controlled specific endonuclease and nuclease(51, 52). Mini-SimCells (size 100-400 nm) are generated through asymmetric division of bacteria with a *minD* gene deletion(53–55). Both SimCells and mini-SimCells can be produced from various bacteria strains(52, 56) and engineered to carry designed DNA for predefined functions(49, 50) In this study, *E. coli* BL21 (DE3) and its ΔminD mutant were used as the chassis to generate SimCells and mini-SimCells, respectively. Both SimCells and mini-SimCells cannot replicate but retain the machinery of cells (e.g. transcription and translation). They are highly controllable and easy to produce from engineered parental bacterial cells(49, 50) In this study, the escape frequency of SimCells is below 10⁻⁸, meeting the criteria of the NIH guidelines for clinical recombinant microorganisms(53). Mini-SimCells are essentially engineered minicells containing designed DNA. Clinical therapy based on minicells has been validated in both dogs and humans (phase I clinical trial), showing significant promise in safety and efficacy(54). Remarkably, minicell-based therapy has recently been granted “Fast-Track” status by the FDA(58). Hence, SimCell and mini-SimCell platforms not only meet safety standards but also show great potential in clinical applications, such as pathogen and cancer treatment(49, 54, 59, 60).

Our strategy for antimicrobial therapy is engineering SimCells and mini-SimCells as smart ‘bio-particles’ to selectively eradicate pathogens, while sparing non-target bacteria (Figure 1). We engineered surface-displayed nanobodies on the outer membrane of SimCells and mini-SimCells using a modular surface display system(61), enabling selective recognition and binding to antigens on target pathogens. This nanobody-antigen interaction not only increases specificity, but also enhances antimicrobial effects that rely on close cell-cell proximity (Figure 1). Such mechanisms include ‘nano-needle’ killing via type VI secretion system (T6SS)(62, 63) and localised release of high dose of antimicrobial compounds. Specifically, SimCells and mini-SimCells engineered with a modularised T6SS are supposed to deliver toxic effectors(64) to the cytoplasm of other bacteria in a contact-dependent manner(65–67). Another mechanism is to locally generate a high concentration of H2O2 via catalysis of aspirin by NahG salicylate hydroxylase enzyme(68–72).

**Figure 1.**
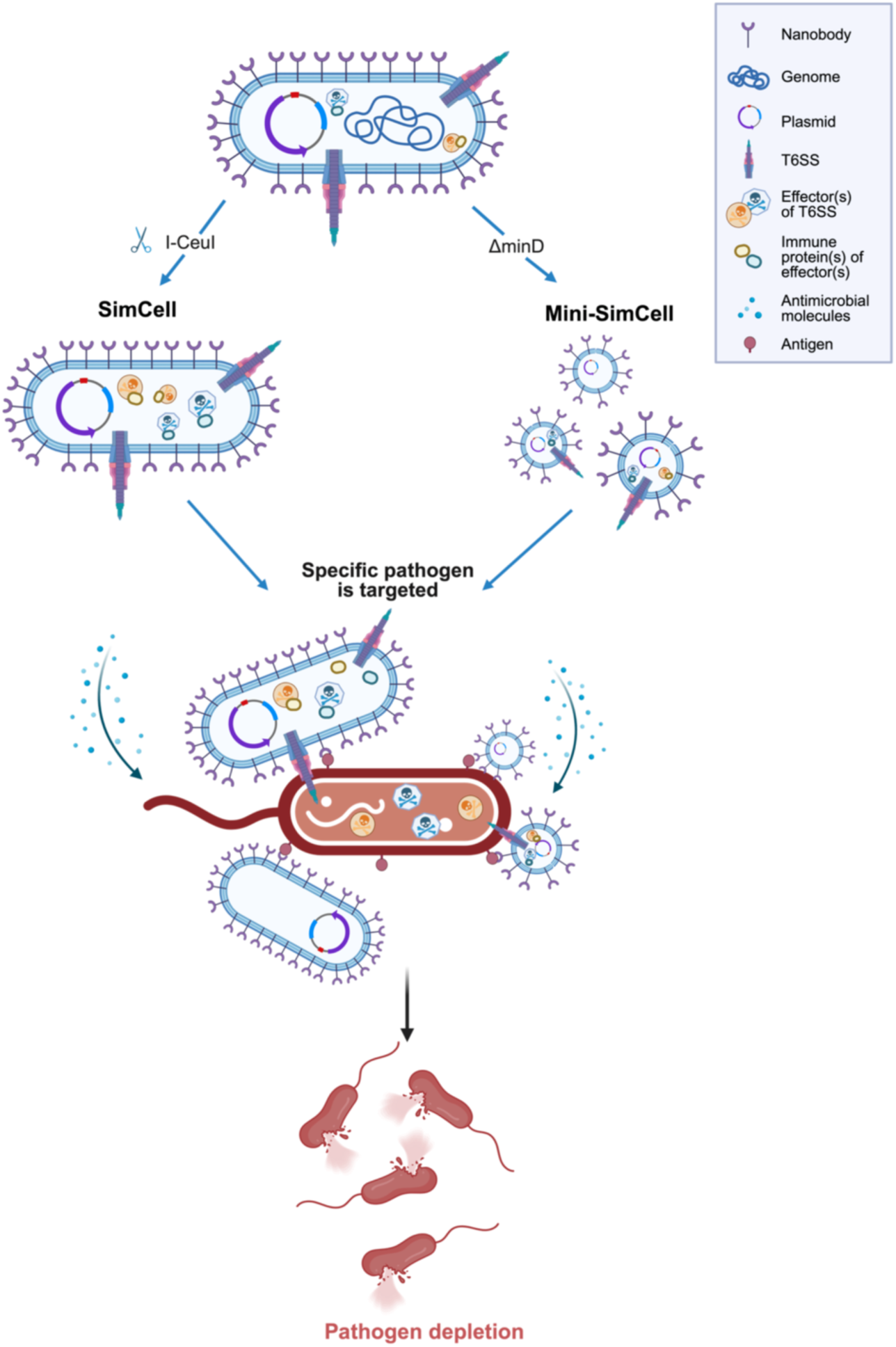
Design of the SimCell and mini-SimCell platform for treating bacterial AMR. Surface-displayed nanobodies are engineered to the outer membrane of SimCell and mini-SimCell using a modular surface display system, to enable antigen recognition and selectively binding to target pathogens. This nanobody-antigen binding is expected to increase the specificity, shorten the distance, and enhance the efficiency of contact-dependent or enhanced inhibition mechanisms including toxic effector protein delivery through modularised T6SS and localised antimicrobial molecules generation.

In this study, we engineered a customisable SimCell and mini-SimCell platform that combined both mechanisms to deliver both rapid and sustained antimicrobial activity. Antimicrobial efficacy is significantly enhanced by the specific binding of SimCells and mini-SimCells to target bacteria via nanobody-antigen interactions, enabling immediate T6SS-mediated cytotoxicity followed by prolonged hydrogen-peroxide release. For proof of concept, a single safe dose of mini-SimCells killed 94% of the target bacteria within 24 hours, and 99% after 48 hours. In mixed microbial communities, four sequential doses achieved a 10³-fold reduction of the target bacteria while leaving non-target species largely unaffected. Engineered with native-antigen targeting nanobody, reprogrammed dual mechanism mini-SimCells successfully eliminated 97% of the target multi-drug resistant pathogen, *E. coli* ST131, within 24 hrs. These findings highlight the potential of SimCells and mini-SimCells for antimicrobial therapy and set a foundation for the development of clinically safe, custom-designable biotherapeutics to combat the escalating threat of antimicrobial resistance.

## Results

### Surface-displayed Nanobody in E. coli Enables Selective Binding

First, the nanobodies and T6SS were characterised with the parental chassis strains before moving on to preparing SimCells and mini-SimCells. *E. coli* BL21(DE3) and its ΔminD mutant were used as the chassis to produce both SimCells and mini-SimCells. The smooth surface without flagella and fimbriae of the B lineage of *E coli* BL21(DE3) minimises interference with the nanobody-antigen interactions. It has been previously reported that an N-terminus-fusion modular system can display nanobodies on the cell’s outer membrane with precise folding and efficient localisation(61). As a proof of concept, we employed a well-characterised antigen-nanobody pair, EPEA (Ag) and anti-EPEA (Nb)(73), to demonstrate the target-specific antimicrobial effect of SimCell therapy (Figure 1).

Surface-displayed nanobody expression and its antigen binding were verified at both macroscopic and microscopic levels. Strains expressing either antigen (Ag) or nanobody (Nb), as well as a null strain, were induced overnight with 100 ng/mL anhydrotetracycline (ATc), and labelled with superfolder GFP (sfGFP) (green) and mRFP (red), respectively. Stationary-phase cultures were normalised to the same optical density (OD600), then left unmixed or mixed in a 1:1 ratio (Ag+Nb and Ag+Null, with the latter as a control without adhesion construct). After 12 hrs standing still, significant cell-cell aggregation was observed in cultures with adhesion pairs, while no such phenomena were seen in unmixed or control groups (Figure 2a). From these, 10 μL cultures were collected from the bottom of both adhesion and non-adhesion groups, and performed the same dilution until single cells were loosely distributed in most views under the fluorescent microscope. Cells with adhesion pairs formed mesh-like-pattern aggregates (Figure 2a). The aggregation was then quantified by measuring the OD600 from the upper 25% portion of the cultures (Supplementary Figure S1a). A significant OD600 reduction (>85%) occurred for the adhesion group (Ag+Nb) (Figure 2b). In the time-course experiment, supernatant density was measured every 30 min for 3.5 hrs. The OD600 reduced by more than 50% for the adhesion group within 1 hr (Figure 2c), indicating that Ag-Nb binding interaction is a rapid process.

**Figure 2.**
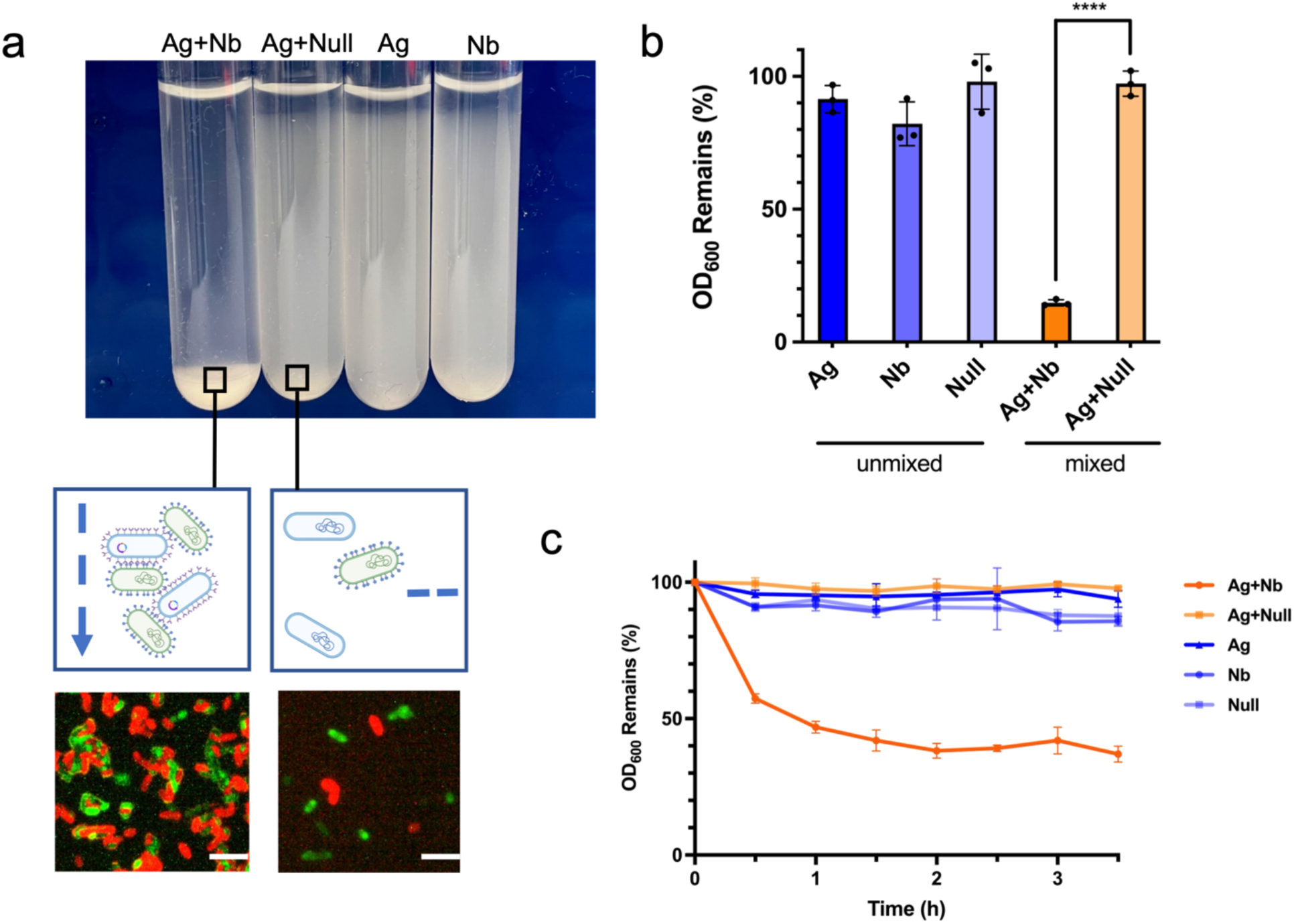
**Surface-displayed nanobody in engineered *E. coli* enables selective cell-cell binding via nanobody-antigen interactions**. **(a)** Binding of Ag and Nb cells in a 1:1 ratio leads to both detectable macroscopic and microscopic aggregation compared to unmixed and non-adhesion (Ag+Null) conditions. Red: mRFP-labelled Nb/Null cells. Green: sfGFP-labelled Ag cells. Scale bar, 10 μm. **(b)** Ag-Nb aggregating cell mixture shows significant settling compared to unmixed and non-adhesion (Ag-Null) conditions after 12 hrs. **(c)** Time courses of macroscopic aggregation show significant settling within ∼ 30 min. All cell cultures were mixed at time 0 and allowed to settle. OD600 from top 25% supernatant was continuously measured every 30 min for 3.5 hrs. For **(b)** and **(c)**, all cultures were normalised to the same initial OD600 and mixed in 1:1 v/v ratio. N = 3 replicates, Error bars, ±1 SD, ****p < 0.0001 according to a 2-tailed paired t-test.

The results suggest that the anti-EPEA Nb should be correctly folded and surface-displayed on the parental chassis strains. To further test nanobody stability, Nb-expressing strains were stored at either room temperature (RT), 20 °C or 4°C for 1, 5, and 10 days, then mixed with Ag-expressing *E. coli*, and macroscopic aggregation was monitored for 3 hrs (Supplementary Figure S1a). There were no significant cell pellets observed in all the unmixed groups, indicating cell integrity and stability. Despite the extended time required to attain comparable aggregation intensity, nanobody activity was preserved in cells stored at both temperatures after 10 days (Supplementary Figure S1b).

### Nanobody–Antigen Pairs Enable Selective T6SS Killing of Target Cells

A plasmid containing complete gene cluster encoding T6SS from *Aeromonas dhakensis*(64), has been introduced to parental chassis strains (Supplementary Table S2). The plasmid contains a sfGFP-fused T6SS gene cluster controlled by either a native constitutive promoter (pT6S_NP) or a tetracycline-inducible promoter (pT6S_Tet), as well as inactive controls (pT6S_N3-NP and pT6S_N3-Tet), which contain a noncontractile 3-amino-acid insertion in the subunits forming contractile outer sheath. Fluorescence microscopy analysis demonstrated the formation of contractile tubular structures, confirming T6SS activity (Figure 3a). *E. coli* BL21(DE3) engineered with T6SS also proved significant killing effect in contact-promoted (solid) condition (Figure S2), confirming T6SS activity. Over 97% of constitutive T6SS-expressing cells exhibited active sfGFP after 96-hour (Figure S2b) and maintained significant out-competitive capability against the prey cells (Figure S2c). These results indicate stable and robust expression of the T6SS across the cell population.

**Figure 3.**
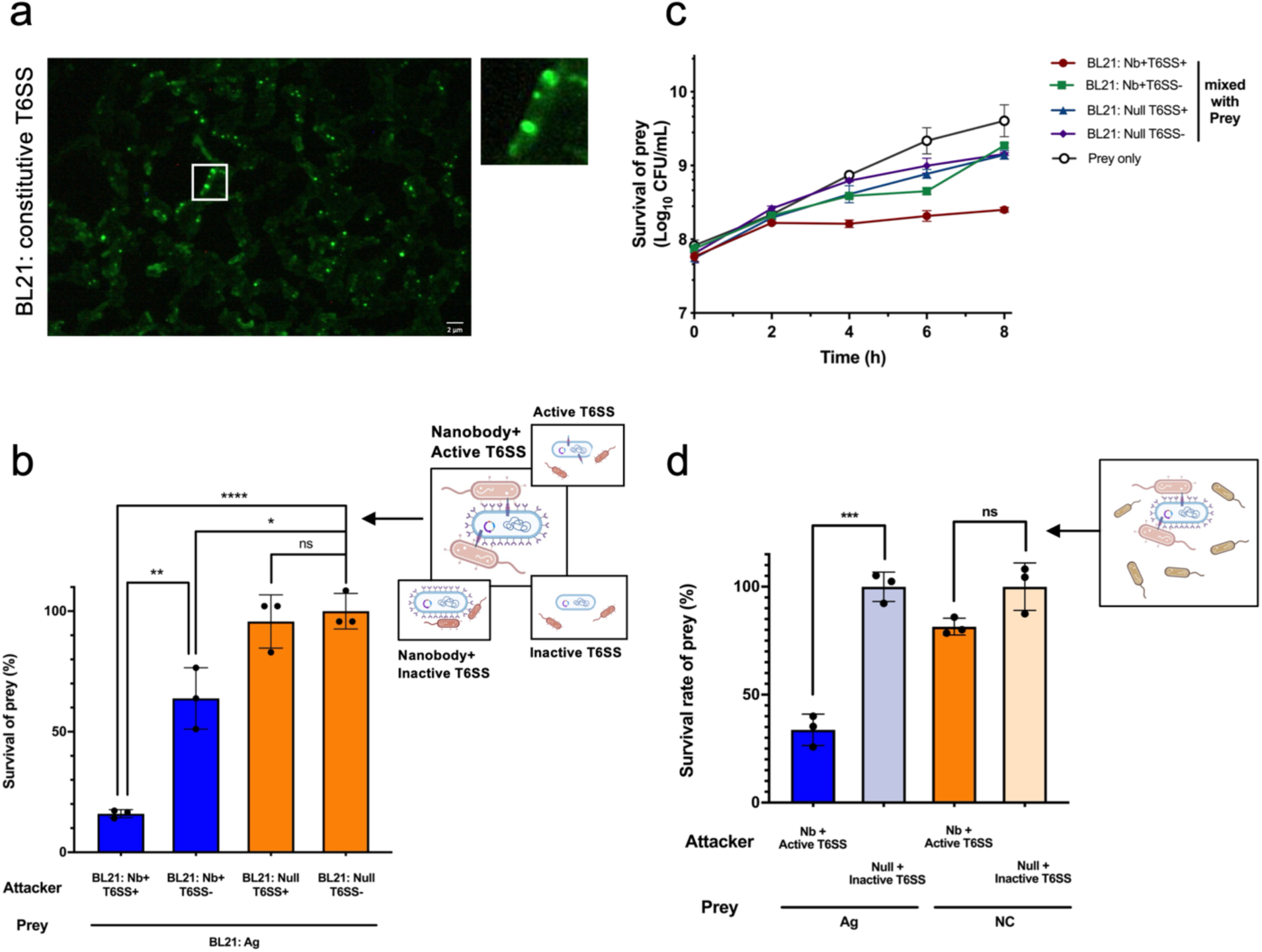
Nanobody-antigen mediated cell-cell adhesion enables selectively depletion of target cells via T6SS in liquid condition. **(a)** Fluorescence microscopic analysis confirms the assembly of T6SS in laboratory *E. coli* strain BL21(DE3). C-terminal sfGFP was fused to VipA for labelling. Localised fluorescence signals were observed. Scale bar, 2 μm. **(b)** Nanobody-antigen binding enables an efficient T6SS-mediated killing in liquid condition. Incubation time = 6h. Attacker : prey v/v ratio = 10:1. In the schematic, target cells with surface antigen and programmed attacker cells (Nb+T6SS+/ Nb+T6SS- / Null T6SS+ / Null T6SS-) are represented. Nb+ = cells with surface-displayed nanobody, Null = cells with no adhesin. T6SS+ = cells with active induced T6SS, T6SS- = cells with inactive induced T6SS. **(c)** Time courses of nanobody-enhanced T6SS killing effect in liquid condition. Significant killing was observed after 2 hours. Incubation time =0, 2, 4, 6, 8 hr. Attacker : prey v/v ratio = 10:1. **(d)** Nb-Ag recognition enables a specific T6SS-mediated killing in liquid condition. In the schematic, programmed attacker cells (Nb+T6SS+) and Prey 1 (Ag), Ag-expressed *E. coli* BL21, prey 2 (NC), *E. coli* DH10B are represented. Incubation time = 6 hr. Attacker : prey 1: prey 2 v/v ratio = 10:1:1. For **(b)** to **(d)**, all cultures were normalised to the same initial OD600. N = 3 replicates, Error bars, ±1 SD, *p < 0.1, **p < 0.01, ***p < 0.001, ****p < 0.0001 according to a 2-tailed paired t-test. Survival rate was calculated as survival cells/ total prey cells.

As the function of T6SS requires close cell-cell proximity and sustained interactions(74), we hypothesised that nanobody-mediated selective binding could promote T6SS^+^ attacker cells to efficiently kill the targeted prey cells in a mixed population. Supplementary Figure S3a show that Nb-Ag interaction enhanced T6SS-mediated killing in solid conditions. In comparison to the contact-promoting killing on the solid surface, well-mixed suspension conditions are representative of clinical scenarios with planktonic pathogens. The results demonstrate that in a fluid environment, the attacker cells with both nanobody and active T6SS have a significant killing effect, with more than 84% of prey cells eliminated compared to the negative control group. However, the attacker cells with active T6SS (T6SS^+^) without nanobodies were unable to eliminate prey cells (Figure 3b). Interestingly, an inhibitory effect was also observed in the Nb-binding-only group (Figure 3b), which may be attributed to the disruption of carbon-source uptake, motility, and membrane integrity in prey cells due to cell-cell aggregation(75). The synergistic functionality of both binding and T6SS^+^ was markedly more pronounced. In the time-course experiment, a significant inhibition effect was detected within 4 hours and continued to increase over time (Figure 3c).

To verify that the surviving prey cells had not acquired resistance to T6SS-mediated killing, we tested various multiplicities of infection (MOI). Increased killing efficiency was observed with higher MOI, suggesting that the inability to eliminate 100% of the prey cells could be due to exhaustion of T6SS activity rather than prey resistance (Supplementary Figure S3b). This limitation could potentially be addressed by increasing the attacker-to-prey ratio, regeneration of T6SS activity or administering multiple doses.

To evaluate whether the nanobody display confers target specificity to T6SS^+^ attacker cells, we co-cultured the attacker cells with a mixed prey population of Ag-positive *E. coli* BL21 (Ag^+^) and Ag-negative *E coli* DH10B (Ag^-^). Selective killing should occur only when (i) the nanobody matched the paired antigen and (ii) attacker cells possessed an active T6SS. After 6 hours of co-incubation, more than 50% of Ag^+^ cells were eliminated, whereas the Ag^-^ population showed no significant loss (Figure 3d and Supplementary S3c). The results confirm that nanobody-antigen interactions enabled precise and T6SS-dependent elimination of the targeted prey bacteria, while sparing non-target bacteria in a mixed microbial community.

### T6SS Mediated Bacterial Killing in Reprogrammed SimCells

Building on the demonstrated nanobody-antigen specificity and T6SS-mediated killing, we next generated SimCells from our chassis strains. We previously reported the development and characterisation of SimCells as a safe, reprogrammable, and customisable chassis for synthetic biology(52), especially beneficial for biomedical applications such as cancer therapy(49). The chromosome-free SimCells were produced by DNA cleavage at a conserved 19-bp recognition sequence (contained within a 26-bp fragment) in the 23S rRNA subunit using inducible expression of I-CeuI endonuclease, after which the bacterial native nuclease RecBCD initiated chromosomal degradation(51).

Plasmids containing the I-CeuI endonuclease, the surface-displayed nanobody and the T6SS machinery were introduced into the *E. coli* BL21 (DE3) chassis to produce and reprogram SimCells (Supplementary Table S2). After the culture reached the early log phase (OD600 ∼ 0.4), 1 μM of crystal violet was added to induce the I-CeuI expression. Growth arrest was observed within 1 hour for SimCell (+) groups, resulting in a stable OD600 plateau maintained for over 35 hrs of continuous culturing, while the uninduced or control groups continued normal growth until reaching the stationary phase. For further verification, 5 μL of post-35-hr cultures were plated on LB agar, and no single colony formation was detected after 48 hrs incubation for induced SimCells (Figure 4a). Additionally, as a second validation, we stained cells with SYTO 9, a DNA-binding fluorescent dye. As expected, no significant fluorescence was observed in SimCells compared to the control group (parental cells) (Supplementary Figure S4a). Single-cell Raman analysis revealed the absence of the Raman band at 785 cm^-1^ in SimCells (Supplementary Figure S4b). Both above evidence confirm the removal of genome DNA. Finally, plating 200 μL of SimCell culture at OD600 = 2.0 (∼10⁸ cells) yielded no single colony formation after 24 hours of incubation (Supplementary Figure S4c). It confirms that the resulting SimCells had high purity, with an escape frequency of less than 10^-8^, meeting the criteria of NIH guidelines for recombinant microorganisms(57).

**Figure 4.**
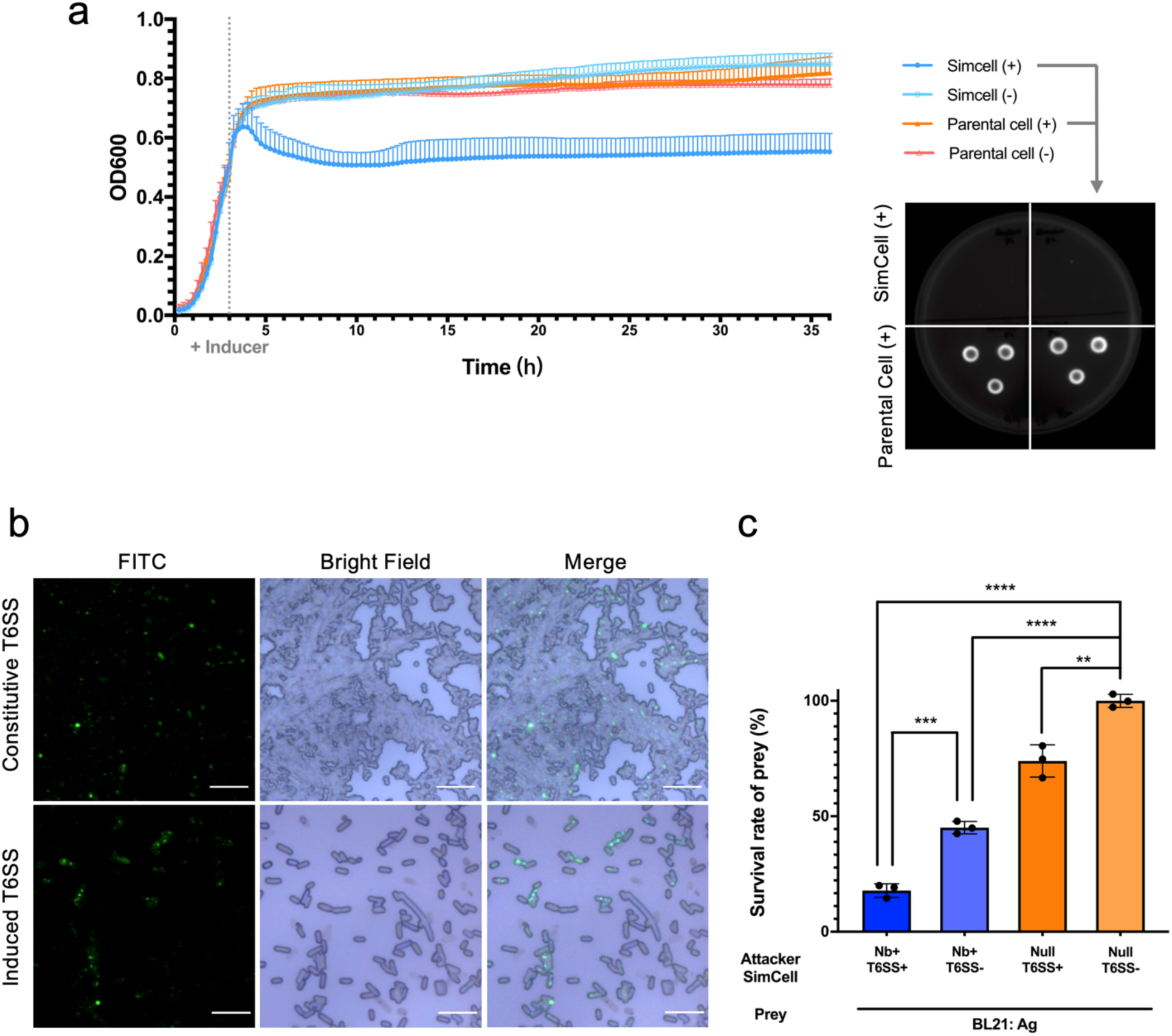
T6SS-mediated killing in reprogrammed SimCells. **(a)** SimCell conversion and purity check. Growth assays confirm the successful SimCell conversion from the engineered parent cell. OD600 was continuously measured every 15 min to monitor the growth curve over 35 hours. The grey dot vertical line indicates the time point (3h after initial incubation) of inducer addition (1 μM crystal violet). Growth arrest was observed shortly after inducer addition for induced SimCells. n = 3 replicates, Error bars, ±1 SD. Agar plate culturing confirmed a high efficiency of SimCell conversion. All cultures were plated on LB agar plates and incubated for 24 hours at 37 °C. **(b)** Fluorescence microscopic analysis confirms the assembly of T6SS in *E. coli* strain BL21(DE3)-derived SimCells. Scale bar, 10 μm. **(c)** Nanobody-antigen binding enhances T6SS-mediated killing in SimCells in liquid condition. Incubation time = 6 hr. Attacker : prey v/v ratio = 10:1. N = 3 replicates, Error bars, ±1 SD, **p < 0.01, ***p < 0.001, ****p < 0.0001 according to a 2-tailed paired t-test. Survival rate was calculated as survival cells/ total prey cells.

We then assessed the binding ability and T6SS activity of the reprogrammed SimCells. The expression of both the nanobody and T6SS was induced prior to SimCell conversion to ensure the engineered proteins were fully assembled for operation. As shown in Figure S1c, significant macroscopic aggregation and OD600 reduction (>60%) were detected for the adhesion group (Ag+Nb-SimCell) after 3 hrs standing. Under the microscope, dotted GFP signals were identified as a characteristic mark for T6SS activity in most SimCells (Figure 4b). SimCells with constitutive and tetracycline-induced T6SS were generated from the cultures with the same cell density. A higher yield of functional SimCells was usually obtained for the constitutive T6SS group. The constitutive T6SS plasmid (pT6S_NP/pT6S_N3-NP) allowed the parental cells to synthesise and assemble an active secretion system before chromosome removal, reducing metabolic burden and yielding more functional T6SS SimCells than an inducible plasmid (pT6S_Tet/pT6S_N3-Tet). Hence, SimCells carrying the constitutive T6SS plasmid were used in subsequent experiments. On agar media, the T6SS activity showed significant killing effect on prey cells within 3 hrs (Supplementary Figure S4d). Nonetheless, SimCell killing efficiency was lower than that of the parental bacteria, likely due to their limited residual energy reserves.

To validate nanobody-mediated, T6SS-dependent killing by SimCells, we co-cultured antigen-positive (Ag⁺) prey with attacker SimCells carrying different combinations of nanobody (Nb⁺ or Nb⁻) and T6SS (T6SS⁺ or T6SS⁻). After 6 h in minimal medium, Nb⁺T6SS⁺ SimCells eliminated more than 85 % of Ag⁺ cells. In contrast, prey survival was roughly 50 % with nanobody binding alone (Nb⁺T6SS⁻), and moderate inhibition was observed with non-binding T6SS⁺ SimCells (Nb⁻T6SS⁺) (Figure 4c). These results confirm that SimCells retain measurable T6SS activity in suspension and demonstrate that specific nanobody binding is critical for maximising T6SS-mediated killing in liquid culture.

### T6SS Mediated Killing in Reprogrammed Mini-SimCells

Following validation of the nanobody-directed specific T6SS killing in SimCells, we subsequently established design feasibility in the mini-SimCell platform. Reprogrammed mini-SimCells were generated from the aberrant cell division of the ΔminD mutant of engineered parental cells. The *E. coli* BL21(DE3) ΔminD strain was previously developed and verified in our lab(49). Mini-SimCells were purified by gradient centrifugation to separate them from parental cells. The formation of T6SS^+^ mini-SimCells was confirmed using fluorescence microscope (Figure 5a) and scanning electron microscope (SEM) (Figure 5b). In addition, ZetaView analysis was used to characterise the size distribution of the mini-SimCells, where mini-SimCells with T6SS (T6SS mini-SimCells) were identified by sfGFP-labelling. Approximately 91% of mini-SimCells derived from parental cells (BL21 mini-SimCells) fell within the 100-300 nm size range, while over 33% of the Nb- and T6SS-expressing mini-SimCells (T6SS mini-SimCells) were larger than 300 nm (Figure 5b). This finding was consistent with the SEM results, suggesting T6SS mini-SimCells were measured larger, likely due to the surface display of Nb and the assembly of the T6SS machinery, as the tail length of the T6SS ‘nano-needle’ is dictated by the size of the bacterial cell(76). ZetaView concentration analysis of the mini-SimCells revealed approximately 2.20 × 10^10^ T6SS mini-SimCells within a total population of 3.56 × 10^11^ reprogrammed mini-SimCells purified from the original 100 mL culture (Figure 5b). This represents a high production yield, indicating promising potential for future large-scale manufacturing applications.

**Figure 5.**
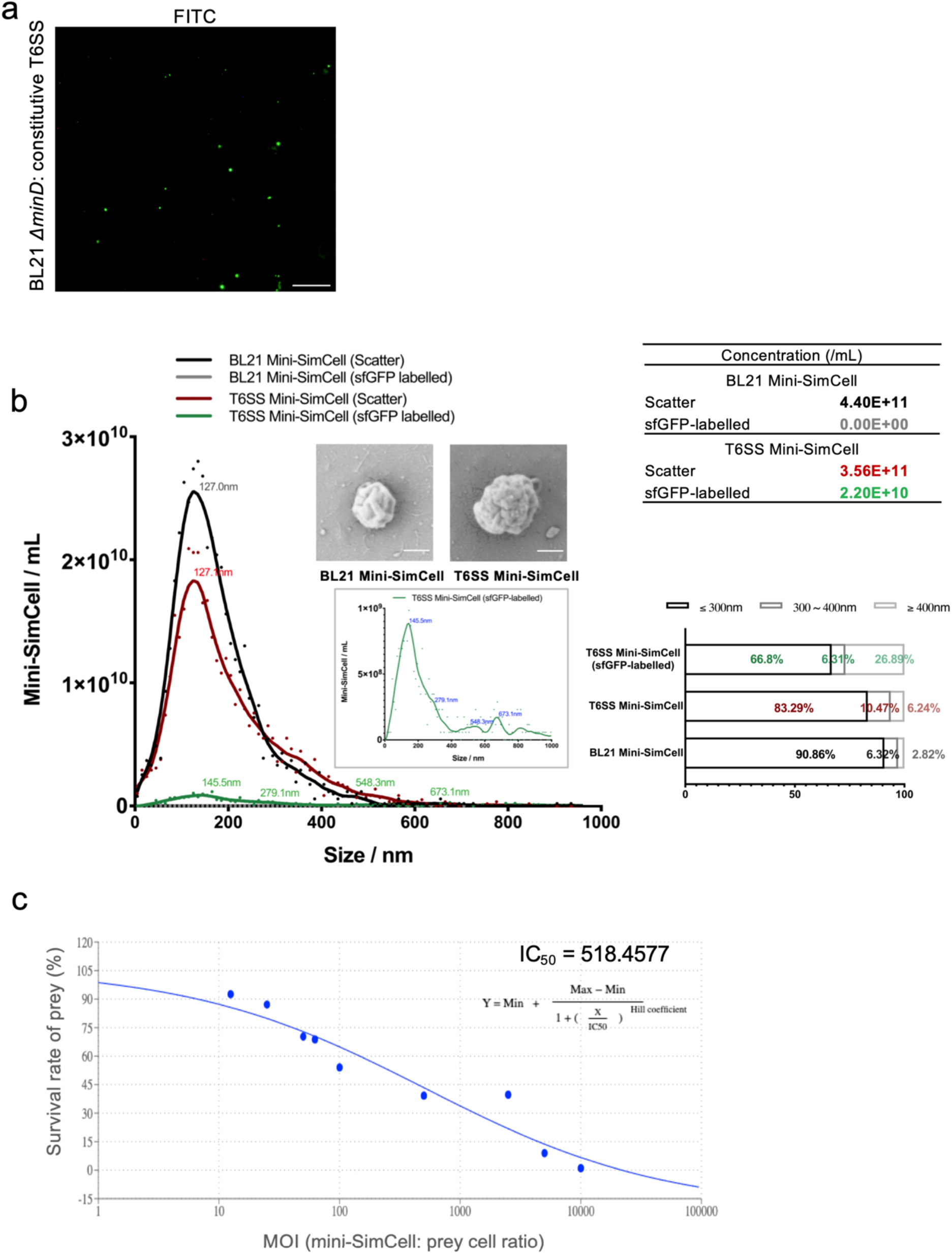
T6SS-mediated killing in mini-SimCells. **(a)** Fluorescence microscopic analysis confirms the assembly of T6SS in *E. coli* BL21(DE3)-derived mini-SimCells. Scale bar, 10 μm. **(b)** Size distribution characterisation of mini-SimCells derived from BL21 parental cells with/without T6SS (T6SS/BL21 mini-SimCells). ZetaView was used for the characterisation of mini-SimCell size distribution and concentration. Scanning electron microscope images of mini-SimCells derived from BL21 parental cells with/without T6SS (T6SS/BL21 mini-SimCells). Magnification, 50,000X. Scale bar, 200nm. **(c)** The inhibitory concentration curve of T6SS-assembled mini-SimCells in liquid condition. A half-maximal inhibitory concentration (IC50) is around 518.5 T6SS-mini-SimCells per prey cell. N = 3 replicates. Survival rate was calculated as survival cells/ total prey cells.

T6SS activity in mini-SimCells was verified on solid media (Supplementary Figure S5a), and nanobodies enabled the T6SS to efficiently kill prey cells also in liquid media. We tested the limit of killing (LOK) for Nb-T6SS mini-SimCells using a dose of 1 × 10^10^ Nb-T6SS mini-SimCells, which has been approved as safe in previous human trials(77). Experiments were conducted at MOI of 62.5, 100, 500, 2500, 5000, and 10,000. As shown in Supplementary Figure S5b, significant inhibition was observed across all groups, with a marked increase in killing efficiency at higher MOI. By fitting an inhibitory concentration curve, the half-maximal inhibitory concentration (IC50) of MOI of Nb-T6SS mini-SimCells was determined to be approximately 518.5 (Figure 5c). The results demonstrate the efficient prey-killing capability of our reprogrammed therapeutic mini-SimCells.

### Localised H2O2 Generation from Parental Cells and (Mini-) SimCells

In addition to the T6SS system, close contact between attacker and prey cells also allows local delivery of high concentrations of antimicrobial compounds around the targeted cells. To exploit this, we introduced a constitutively expressed salicylate hydroxylase (NahG) into our system (Figure S7a), which catalyses the conversion of acetylsalicylic acid (aspirin) into catechol(71, 72) (Figure 6a). Catechol has a broad-spectrum antimicrobial activity(68–70) by generating hydrogen peroxide (H2O2) through autooxidation processes (Supplementary Figure S6a,b,c), during which catechol polymerises to form cross-linked polymers without external catalysts(78–81) (Figure 6a). When 800 μM aspirin (a non-toxic dose for humans(82)) was added to the parental cell and SimCell cultures, the filtered supernatants from overnight NahG+ cultures exhibited a dark-brown colour (Supplementary Figure S7b), which is associated with the oxidation products of catechol. The collected supernatants showed a significant inhibitory effect on bacterial cell growth (Supplementary Figure S7b and S7c). These results indicate the generation, permeability and extracellular antimicrobial activity of SimCell-produced catechol and associated production of H2O2.

**Figure 6.**
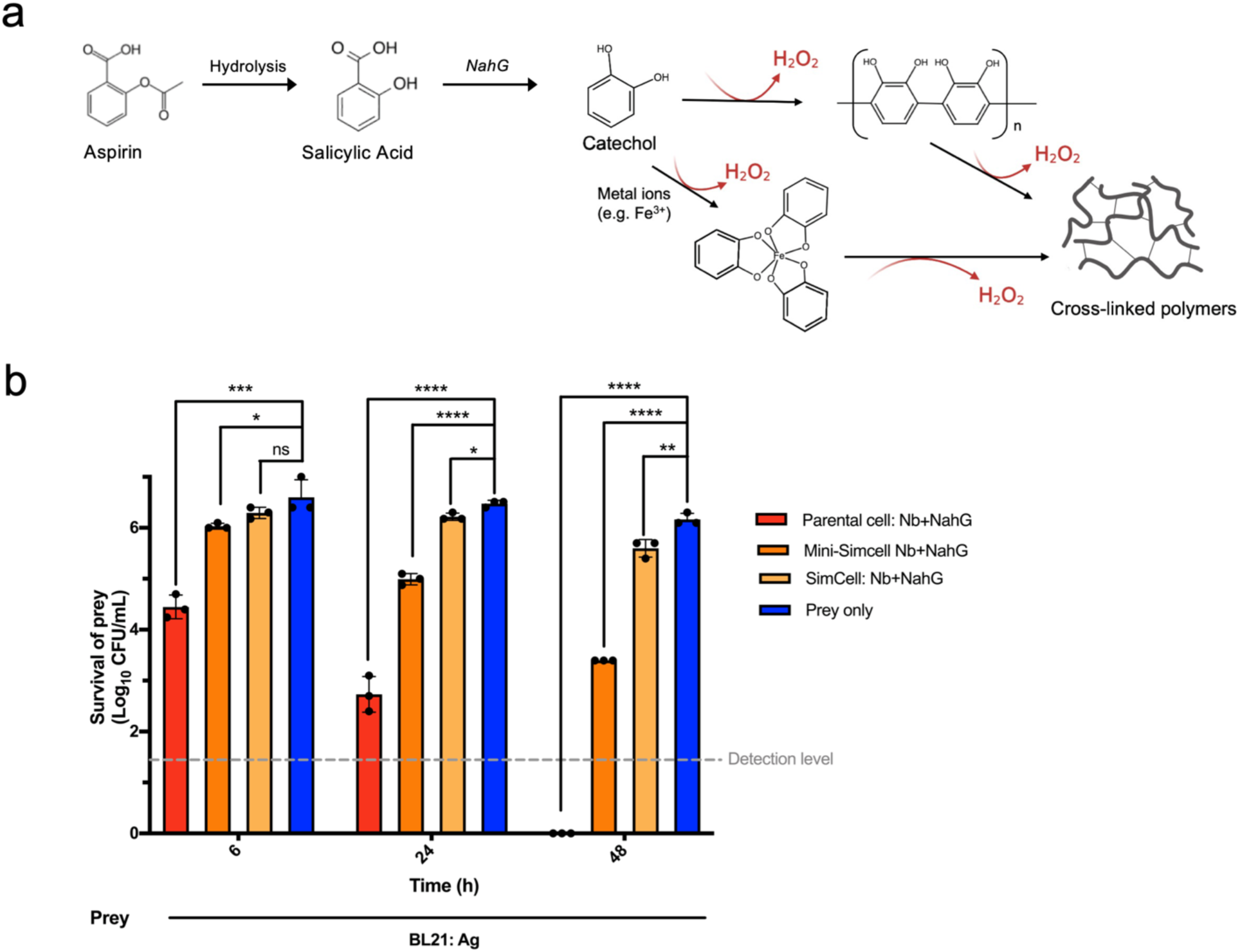
Reprogrammed parental cells and (mini-) SimCells enables localised generation of antimicrobial compound – catechol. **(a)** Schematic illustration of aspirin hydrolysis to salicylic acid, followed by the generation of catechol catalysed by the NahG enzyme. The auto-oxidation of catechol and cross-linking results in the release of H2O2. **(b)** Inhibitory efficiency of localised catechol-producing parental cells / SimCells / Mini-SimCells in liquid condition. Attacker : prey n:n ratio = Parental cells / SimCells / Mini-SimCells 10/100:1,000:1. N = 3 replicates, Error bars, ±1 SD, *p < 0.1, **p < 0.01, ***p < 0.001, ****p < 0.0001 according to a 2-tailed paired t-test.

We then investigated co-expression of nanobody and NahG (Nb+NahG) in parental cells, SimCells and mini-SimCells to eliminate targeted prey cells. The Nb+NahG parental cells, SimCells and mini-SimCells were added to the prey cells at MOI of 10, 100 and 1000, along with 800 μM aspirin. As shown in Figure 6b, significant inhibition was apparent in 6 hrs and pronounced across all experimental groups by 24 hrs. Particularly after 48 hrs, Nb+NahG mini-SimCells eliminated over 99.9% of the prey cells in the presence of a single aspirin dose. These results demonstrate that the NahG reprogrammed SimCells provide a rapid and durable, aspirin-dependent antimicrobial effect.

Notably, NADH is required for NahG catalysis(72, 83). Prior studies suggest that mini-SimCells possess a higher NADH/NAD+ ratio(84, 85), likely due to the reduced electron transport chain activity resulting from the deletion of the *minD* gene and subsequent alterations in the activity of inner membrane enzymes(86, 87). Along with safety and a higher specific surface area (SSA) that enhances catechol penetration, mini-SimCells system is believed to be a promising platform and hence selected for further experiments.

### Dual-mechanism Killing Effect in Mini-SimCells

Building upon these results, we evaluated an integrated mini-SimCell strategy that combines nanobody-mediated binding, T6SS activity and the localised release of an antimicrobial compound. To divide cellular burden and optimise the functionality of two different antimicrobial mechanisms (T6SS facilitated nano-needle and NahG mediated H2O2 local release), we employed a consortium of Nb+ mini-SimCells comprising two specialised subgroups: Attacker1 (mini-SimCell Nb-T6SS) and Attacker2 (mini-SimCell Nb-NahG). Non-engineered mini-SimCells (mini-SimCell Null) were used as negative controls. Antigen-positive prey cells were mixed and co-cultured with various combinations of engineered and control mini-SimCells in the minimal medium. After 6 hrs, 800 μM aspirin was added. The T6SS-mediated attack was triggered immediately upon cell-cell contact, whereas H2O2-based killing began after catechol production from aspirin. Delaying catechol and subsequent H₂O₂ production potentially prevented these toxic metabolites from inhibiting T6SS activity. Prey cell viability was quantified at 6, 24, and 48 hrs without further additions of attacker cells or aspirin.

At an overall MOI of 250:250:1 (Attacker 1:Attacker 2:prey), each engineered mini-SimCell subgroup independently reduced prey viability at different time scales, confirming the efficacy of both antimicrobial modes. When combined, the dual-mechanism consortium achieved the greatest effect, eliminating approximately 90% of prey cells after 48 hrs, demonstrating a synergistic antimicrobial effect with two mechanisms (Figure 7a). We next tested a higher dose dual-mechanism mini-SimCells to treat the target bacteria. Two types of attacker mini-SimCells were mixed with targeted prey cells at MOI of 500:500:1 (Attacker 1:Attacker 2:prey) with the total mini-SimCells of 1×10^10^, the maximum tolerated dose reported in human minicell trials(77). Within the first 6 hrs, the Attacker1 (mini-SimCell engineered with Nb-T6SS) rapidly eliminated over 51% of the target bacteria. Addition of 800 μM aspirin enabled Attacker2 (mini-SimCell engineered with Nb-NahG) to convert aspirin to catechol and generate local H2O2 by polymerising catechol, increasing the killing efficiency to 94.4% ±0.73% at 24 hrs and further reaching 99.3% ±0.29% at 48 hrs (Figure 7b).

**Figure 7.**
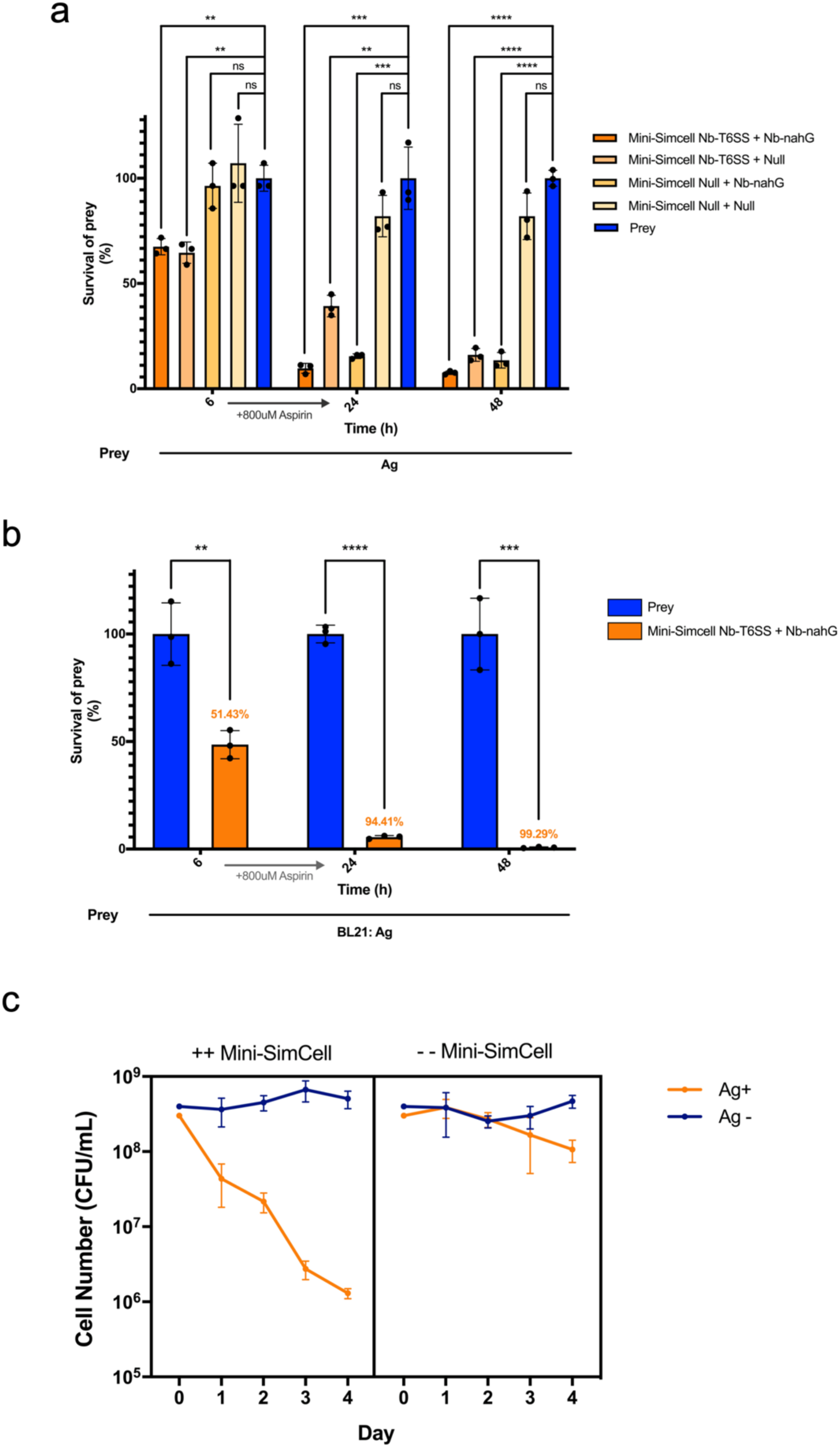
Selective elimination efficiency of the consortium with dual-mechanism mini-SimCells in liquid condition. **(a)** Two different attackers (Attaker1: mini-SimCell Nb-T6SS; Attaker2: mini-SimCell Nb-nahG) were mixed with prey cells in n:n ratio of Attacker1: Attacker2 : prey = 250:250:1, total Attacker cells were 5×10^9^, within the maximum tolerated dose (MTD) reported in the previous human trial. All cultures were incubated at 37 °C and sampled from 6, 24, 48 hr. Aspirin of 800 μM was added after 6 hr. **(b)** Dual-mechanism mini-SimCell inhibition efficiency in MTD for humans. Two different attackers (Attaker1: mini-SimCell Nb-T6SS; Attaker2: mini-SimCell Nb-nahG) were mixed with prey cells in n:n ratio of Attacker1 : Attacker2 : prey = 500:500:1, total Attacker cells were 1×10^10^, the MTD reported in the previous human trial. All cultures were incubated at 37 °C and sampled from 6, 24, 48 hr. Aspirin of 800 μM was added after 6 hr. **(c)** Selective elimination of targeted bacteria population in mixed communities using multi-dose dual-mechanism mini-SimCell consortium. Mixed prey cell populations with or without recognisable antigens (Ag+ or Ag-) were inoculated in n:n ratio of 1:1 for continuous co-culture at 37 °C. 1/50 dilution with fresh medium was conducted every 24 hrs, followed by 8 hrs of recovery growth, then introduction of a dose of mini-SimCell consortium and 16 hrs of co-incubation. For every dose of engineered mini-SimCell (++ Mini-SimCell), two different attackers (Attaker1: mini-SimCell Nb-T6SS; Attaker2: mini-SimCell Nb-nahG) were mixed in n:n ratio of 1:1, total attacker were 1×10^12^ mini-SimCells. For - - Mini-SimCell group, 1×10^12^ of non-engineered mini-SimCell were used. All cultures were incubated at 37 °C and sampled every 24 hrs. Aspirin of 800 μM was added with fresh medium. For **(a)** to **(c)**, N = 3 replicates, Error bars, ±1 SD, **p < 0.01, ***p < 0.001, ****p < 0.0001 according to a 2-tailed paired t-test. Survival rate was calculated as survival cells/ total prey cells. Killing efficiency = 1-survival rate.

To further evaluate the specificity and robustness of the platform in a complex environment, we applied a multi-dose mini-SimCells to selectively target Ag^+^ bacteria population within a mixed microbial community. In this experiment, two distinct types of prey populations: *E. coli* BL21 with or without surface-displayed EPEA antigen (Ag+/Ag-) were continuously co-cultured in a well-mixed condition. To mimic periodic wash-out scenario, the cultures were diluted 1/50 with fresh medium every 24 hrs, allowed to recover for 8 hrs, and then challenged with 10^12^ ++ mini-SimCells (1:1 ratio of Nb-T6SS mini-SimCells : Nb-nahG mini-SimCells) or an equivalent number of the control, which is non-engineered - - mini-SimCells (mini-SimCell without introduced plasmid). After each dose, co-cultures were incubated for an additional 16 hrs to allow mini-SimCell mediated killing to run completely. In the presence of 800 µM aspirin, four sequential doses of mini-SimCells, selective elimination of the Ag^+^ population was observed only in the ++ mini-SimCells treatment group (Figure 7c). Progressive elimination of Ag+ cells occurred following each dual-mechanism mini-SimCell administration, ultimately achieving a 10^3^-fold reduction relative to the initial prey cell levels (Figure 7c). This selective elimination was not observed in the control groups (- - mini-SimCell group), ruling out different growth rates between Ag+ and Ag- populations in the microbial community (Figure 7c). Furthermore, the Ag- population remained stable throughout the treatment experiment, confirming the specificity of the target elimination (Figure 7c). Collectively, these results demonstrate the potent and sustained antimicrobial activity of dual-mechanism mini-SimCells. The combination of rapid T6SS-mediated killing with prolonged NahG-driven H₂O₂ production has effectively and selectively eradicated targeted cells in mixed communities while sparing non-target populations.

### Reprogrammed Mini-SimCells Effectively Targeted and Eliminated E. coli ST131, an AMR Pathogenic Strain

Building on the proof-of-concept platform demonstrating SimCell- and mini-SimCell-based targeted antimicrobial activities in mixed communities, we then evaluate the system against a clinically relevant multidrug-resistant pathogen. *E. coli* ST131 represents a globally dominant AMR strain responsible for a substantial proportion of urinary tract and bloodstream infections(88–91). It raises clinical concern particularly because of its resistance to fluoroquinolones and β-lactam antibiotics due to its extended-spectrum β-lactamases (ESBLs)(89, 92). The WHO has classified ESBL-producing *E. coli*, including ST131, as a high-to-critical priority AMR pathogen, underscoring the urgent need for novel therapeutic strategies targeting this AMR strain (93). Hence *E. coli* ST131 (NCTC 13441) strain was selected for the evaluation of the SimCell efficacy against AMR pathogens.

To achieve specific recognition of *E. coli* ST131, we engineered (mini-)SimCells to express nanobodies targeting outer membrane protein (OmpA). Specifically, we used Nb39, a nanobody exhibiting high binding affinity and specificity for the long OmpA isoform (OmpA-L) in *E. coli* isolates(94). We engineered Nb39 fused with the N-terminal modular system for surface display in the parental strain of mini-SimCells. Stationary-phase cultures of *E. coli* ST131 and Nb39-displaying cells (BL21:Nb39-mRFP) were adjusted to the same OD600, mixed at a 1:1 ratio, and allowed to settle overnight undisturbed. *E. coli* BL21(DE3) strains engineered with surface displayed synthetic antigen (BL21: Ag), EPEA, were used as the unrecognised control and mixed with Nb-39-displaying cells. *E. coli* ST131 and BL21: Ag cells were pre-stained with SYTO 9, while Nb39-displaying cells were genetically labelled with mRFP for fluorescence tracking. Figure 8a shows parental cells (expressing mRFP) with Nb39 surface display bond to *E. coli* ST131 (SYTO9 staining). Significant cell-cell aggregates were observed under fluorescence microscopy (Figure 8a Left), whilst the control showed scattered cell distribution (Figure 8a Right). Aggregation of these cells was mediated by the interactions of the OmpA-L antigen and Nb39 nanobody. Flow cytometry detected single events exhibiting both green and red fluorescence (G+R), designated as “doublets”. Flow cytometry analysis demonstrated that 46.4% of the population exhibited doublet characteristics (Figure 8b Left), whilst the control had 0.0% doublets (Figure 8b Right). It indicates the efficient targeting at *E. coli* ST131 via Nb39-mediated antigen recognition. These results confirm that surface-displayed Nb39 enables specific and robust binding to *E. coli* ST131 through native antigen (OmpA-L) recognition, preparing for the targeted mini-SimCells killing.

**Figure 8.**
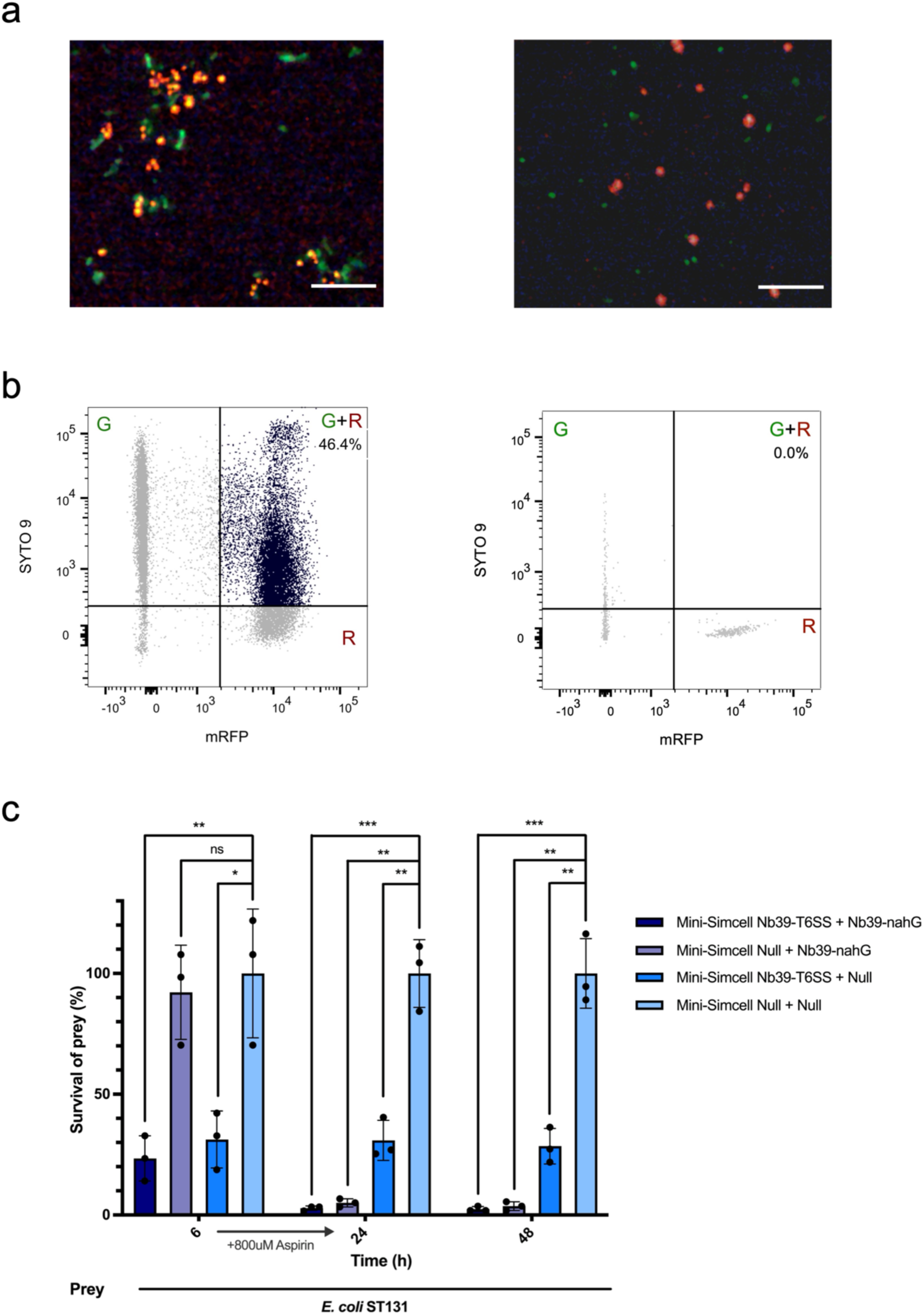
Reprogrammed mini-SimCells to kill a resistant pathogenic *E. coli* strain (*E. coli* ST131) by targeting a native antigen (OmpA-L). (a) Binding assay: *E. coli* ST131 and mini-SimCell parental strain BL21: Nb39_mRFP were mixed 1:1 ratio, leading to significant aggregation due to the Nb39-OmpA-L interaction, under fluorescence microscope (left). BL21: Ag and BL21: Nb39_mRFP cells were mixed with as unbinding negative control (right). Red: BL21: Nb39_mRFP cells. Green: SYTO 9 stained *E. coli* ST131 cells (left), SYTO 9 stained BL21: Ag cells (right). Scale bar, 10 μm. **(b)** Flow cytometry assay: Quantification of Nb39-mediated *E. coli* ST131 binding using a double-color events. Events of doublets formation (G+R) indicates the binding of *E. coli* ST131 and BL21: Nb39_mRFP. BL21: Ag and BL21: Nb39_mRFP mixture was the unbinding negative control. *E. coli* ST131 (left)/BL21: Ag (right) : BL21: Nb39_mRFP cell = 1:1. **(c)** Inhibitory efficiency of the consortium with dual-mechanism mini-SimCells against *E. coli* ST131. Two different attackers (Attaker1: mini-SimCell Nb39-T6SS; Attaker2: mini-SimCell Nb39-nahG) were mixed with prey cells in n/n ratio of Attacker1: Attacker2: prey = 500:500:1, showing elimination effect over 70%, 97% and 97% at 6, 24 and 48 hrs. All cultures were incubated at 37 °C and sampled from 6, 24, 48 hrs. Aspirin of 800 µM was added after 6h. N = 3 replicates, Error bars, ±1 SD, *p < 0.1, **p < 0.01, ***p < 0.001 according to a 2-tailed paired t-test. Survival rate was calculated as survival cells/ total prey cells; killing efficiency = 1-survival rate.

Subsequently, we demonstrate that reprogrammed mini-SimCells achieved efficient elimination of the multidrug-resistant pathogenic strain *E. coli* ST131 (Figure 8c). Nb39-displaying mini-SimCells equipped with either T6SS (mini-SimCell: Nb39-T6SS) or nahG (mini-SimCell: Nb39-nahG) modules were evaluated for their antimicrobial efficacy. *E. coli* ST131 elimination assays were conducted at an overall MOI of 500:500:1 (Attacker 1: Attacker 2: Prey), with a total mini-SimCell number of 1×10¹⁰. Mini-SimCells equipped with nanobody-directed T6SS achieved >70% elimination of *E. coli* ST131 at 6 hrs (Figure 8c). When 800 μM aspirin was supplemented, the converted catechol and locally generated H2O2 further enhanced antimicrobial activity. Hence, the dual-mechanism mini-SimCell consortium increased the elimination efficiency to 97.19% ±0.97% at 24 hrs and 97.58% ±1.05% at 48 hrs (Figure 8c). In contrast, *E. coli* ST131 exhibited persistent resistance to broad-spectrum β-lactam antibiotics, with no growth inhibition observed for benzylpenicillin (PenG) and cefotaxime even at 100 μg/mL (supplementary Figure S8a and S8b). These results demonstrate that the nanobody-targeted, dual-mechanism mini-SimCell platform can overcome conventional antibiotic resistance mechanisms to achieve potent, specific elimination of clinically relevant multidrug-resistant pathogens.

## Discussion

In this study, we introduce a novel SimCell and mini-SimCell platform with antimicrobial properties. Through nanobody-antigen binding, the reprogrammed SimCells and mini-SimCells are designed to selectively eliminate target cells via T6SS-mediated killing within 4-6 hrs and local conversion of aspirin to catechol, which then continuously generates high-concentration of H2O2 over several days. This antimicrobial strategy provides both immediate and sustained antimicrobial effects, offering the potential for controlling acute bacterial infection and preventing reoccurrence. For proof-of-concept, our results demonstrate that the mini-SimCell consortium can eliminate 94.4% of target prey cells in 24 hrs, and 99.3% in 48 hrs with a single dose of mini-SimCells at MOI of 1000:1. Our results also demonstrate the selective pathogen depletion up to 10^3^-fold within the mixed microbial communities using multi-dose mini-SimCell administration. This observation further validates the specific antimicrobial efficacy of the platform. To demonstrate the anti-AMR capacity and clinical application potential of our reprogrammed (mini-)SimCell-based therapeutic platform, we adapted the system to target the clinically relevant, multidrug-resistant pathogenic strain *E. coli* ST131. Equipped with a native OmpA recognising nanobody (Nb39), our reprogrammed mini-SimCells achieved 97.19% target pathogen elimination within 24 hours, overcoming conventional antibiotic resistance mechanisms. This success against a high-to-critical priority AMR pathogen demonstrates the translational potential of our platform for addressing urgent clinical challenges posed by multidrug-resistant infections. The combination of antibacterial efficacy and selectivity represents a significant advancement over broad-spectrum antimicrobial approaches, positioning this platform as a promising next-generation therapeutic strategy for combating antimicrobial resistance.

We vision this platform as a new approach for developing clinically safe, custom-designable biotherapeutic products against multi-drug-resistant pathogens. The key attributes and advantages of this platform include the following aspects: 1) Biosafety. SimCells and mini-SimCells met the criteria of NIH-recommended escape frequency threshold - one live cell from 10^8^ SimCells (57). Minicells derived from *Salmonella typhimurium,* which are similar to *E. coli* derived mini-Simcells, have been granted US FDA approval for human trial (https://engeneic.com/), and proven safe without adverse effect in human phase I clinical trial (95). To further enhance biosafety, our platform will be extended to use SimCells and mini-SimCells derived from LPS-free *E. coli* BL21(DE3) chassis (ClearColi)(96).

2) Antimicrobial specificity and efficacy. SimCell and mini-SimCell chassis are highly controllable while maintaining robust bioactivity against target bacteria. The precise targeting helps to minimise unintended effects on non-targeted microbes, enhancing the therapeutic efficacy. 3) Modularity and customisability. This modular platform features a versatile architecture that allows for easy redesign in a “plug-and-play” manner. Surface-displayed nanobodies can be systematically reconfigured to target a specific new pathogen, enhancing the adaptability of the platform. Additionally, T6SS effectors can be engineered and seamlessly integrated into the existing modules. The enzyme-catalysis mechanism for converting pro-drugs to antimicrobial compounds, such as the aspirin-catechol system, can also be redesigned to fit specific therapeutic needs. 4) Scalability. SimCells and mini-SimCells can be easily produced and purified from engineered parent cells, making them suitable for large-scale industrial manufacturing. The streamlined production process, combined with high yield and cost-effective scalability, offers a manufacturing practical solution to address the urgent challenges posed by AMR.

The historic battle between humans and bacterial pathogens has intensified since the “Golden Age” of antibiotic discovery in the 1950s(8). Today, only a limited number of small molecules are currently undergoing clinical trials as potential antibiotic candidates, with nearly half being new beta-lactamase inhibitors used with established beta-lactam antibiotics(97). Recent advances in AI and machine learning (ML) have accelerated antibiotic discovery and design. A landmark study employed a deep neural network to identify a novel broad-spectrum antibiotic, halicin(22), followed by works utilising machine learning models to discover new antibiotics against drug-resistant *Acinetobacter baumannii*(19, 20). A recent work for the first time *de novo* designed structurally novel antibiotics for drug-resistant gonorrhoea and MRSA via a generative deep learning approach(23). Despite these advances, the “black box” nature of some AI models can obscure the rationale behind predictions and limit the exploration of chemical spaces. Hence, researchers are now actively working on interpretable AI models(21). Additionally, validation of these in silico predictions and data collection for model training still requires extensive experimental tests.

Traditional chemical-based antibiotics face an evolutionary paradox: while effectively eliminating susceptible bacteria, they also promote the survival and spread of resistant variants, inadvertently advancing AMR with each treated infection(98). In response, alternative antimicrobials have been developed, including antimicrobial peptides, bacteriophage, and monoclonal antibodies. Each approach faces its challenges, such as biosafety, stability and the potential risk of rapid resistance development (Supplementary Table S1). Several studies have reported the antimicrobial effects of bacteria engineered to neutralise virulence(99, 100) or eradicate pathogens. To selectively inhibit the target bacteria, toxic cargos were delivered by conditional lysis-based release(101, 102) or cytoplasm-translocation via conjugation(103–105). Nonetheless, balancing high efficacy with the management of side effects (e.g. endotoxin release during lysis) remains a challenge for these approaches. In contrast, the highly controllable and safe SimCells and mini-SimCells are reprogrammed to selectively target pathogens through surface-displayed nanobodies and efficiently eliminate the target cells, using contact-dependent antimicrobial strategies.

This study established a modular antimicrobial platform as proof of concept, and further successfully demonstrates its adaptability using a native-antigen-targeting nanobody against a multi-drug resistant pathogen from a clinical isolate. In real world clinical practice, the surface antigens of AMR pathogens could change due to mutations and environmental factors, potentially affecting epitope accessibility. The modularised design of mini-SimCell platform can be rapidly adapted to equip new nanobodies in response to the changes of surface antigens in AMR pathogens. We expect to further expand nanobody reservoirs targeting antigens from various multi-drug-resistant pathogens and resembling diverse clinical scenarios. The current landscape of nanobody discovery relies on immunity-based approaches(106), which restricts diversity and availability. Emerging strategies, such as the combination of bioinformatics-driven antigen mining and high-throughput library(107, 108), AI enabled *de novo* protein design(109, 110) and deep learning models(111), offer promising approaches to overcome the limitations and accelerate the expansion of the nanobody repertoire for next generation of precise antimicrobials. The dual-mechanism platform in this study broadly targets evolutionarily conserved cellular processes, via T6SS ‘nano-needle’ spearing and H2O2 toxicity, minimising the likelihood of resistance development. However, antigen mutations could possibly enable the prey cell to escape from being recognised and targeted. Hence, the development of multiple nanobodies that target different antigens of AMR pathogens could be a potential solution.

T6SS represents a worth-noting strategy in this study. Building on previous research that uses native T6SS as controllable weapons against target pathogens(112, 113), our work advances the field by heterologously expressing and assembling functional T6SS in SimCell and mini-SimCell chassis, enhancing T6SS-based biotherapeutic applications. The natural diversity of T6SS’s effectors and targets provides a foundation for expanding the functionality of engineered T6SS in this SimCell platform to antimicrobial-resistant Gram-positive bacteria and fungi(114). Engineered T6SS could also deliver various protein payloads, such as Cas9(115) and Cre recombinase(116), directly to the cytoplasm of target pathogens. SimCell and mini-SimCell chassis also minimise environment disruption *in situ*, with the versatility of T6SS opening possibilities for a wide range of biotherapeutic interventions(115) and ecological modification of microbial communities(117).

In conclusion, the reprogrammed antimicrobial SimCell and mini-SimCell platform described in this study offers a novel approach for tackling global bacterial AMR challenges. This work exemplifies the potential of synthetic biology to revolutionise biotherapeutic applications, demonstrating an innovative and effective approach to combat persistent and emerging AMR threats.

## Materials and Methods

### Bacterial strains and growth conditions

All strains used in this study are listed in Supplementary Table S2. *E. coli* DH5α and DH10B were used for cloning and plasmid maintenance. *E. coli* BL21(DE3) was used for the function test and SimCell generation. E. *coli* BL21(DE3) ΔminD was used for mini-SimCell generation. Heat shock and electroporation methods were used for chemical transformation for both strains. Above cells were cultured in Luria−Bertani (LB) media or M9 minimal media with specific nutrient supplements (0.4% glucose, 0.2% casamino acids, 1x trace elements) at 37 °C, 180 rpm. *E. coli* ST131 (NCTC 13441) cells were cultures in M9 minimal media with 0.4% glucose and 1mM EDTA at 37 °C, 180 rpm. Corresponding antibiotics were used for selection: 25 μg/mL chloramphenicol, 50 μg/mL kanamycin, 50 μg/mL carbenicillin, 100 μg/mL streptomycin or 25 μg/mL cefotaxime. Corresponding inducers were used for induction: 100 ng/mL anhydrotetracycline (ATc), 0.2% arabinose (Ara) or 1 μM crystal violet. Strain stocks were stored in 20% glycerol at −80 °C for long-term storage. Agar plates were kept at 4 °C for short-term storage.

### Plasmid construction

All plasmids used in this study are listed in Supplementary Table S2. Plasmids were obtained from lab storage, purchased from Addgene (pDSG287, pDSG289, pDSG291) or as gifts from collaborators. Q5 high-fidelity DNA polymerase (New England Biolabs) was used for DNA fragment amplification. NEBuilder HiFi DNA Assembly (New England Biolabs) was used for plasmid construction. DreamTaq Green PCR Master Mix (Thermo Fisher Scientific) was used for colony PCR. DNA stocks were stored at −20 °C.

### Macroscopic and microscopic aggregation assays

Cultures were grown at 37°C, 180 rpm in 50 mL LB with corresponding antibiotics in a 200 mL flask for 24 hours to ensure stationary phase. All cultures were centrifuged at 3500 rpm for 20 min, resuspended with 1xPBS solution, then adjusted to an initial OD600 of 0.8. Cultures were mixed in a 1:1 v:v ratio (total volume 3mL) in 5 mL clear polypropylene tubes. For macroscopic aggregation, 100 μL of sample was collected from the top 25% supernatant for each measurement. OD600 measurements were read by Synergy 2 microplate reader (BioTek). For microscopic aggregation, 100 μL of the sample was collected from the bottom of the overnight settled mixture and then diluted to an appropriate concentration for fluorescence microscopy using Nikon Ti Eclipse. Fiji was used for image analysis.

### Bacterial Killing Assay

For parental cells, overnight cultures of attacker and prey cells were transferred into fresh LB medium with corresponding antibiotics at a ratio of 1:100 and grown to an OD600 = 1, the inducer was added when OD600 reached 0.6 if needed. Cells were harvested by centrifugation at 1,500 g for 10 min, and the pellets were resuspended in fresh LB or M9 minimal media. For SimCells and mini-SimCells, cell pellets were washed twice with fresh LB or M9 minimal media to fully remove the crystal violet or antibiotics. Cell suspensions were normalised to an initial OD600. Mixtures of attacker and prey cells were then established at intended v:v or n:n ratios, cell number is calculated as(118, 119):

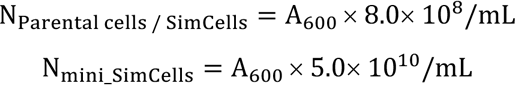

For solid (contact-promoting) condition, triplicates of 5 μL of each mixture were spotted on LB / M9 agar plates (inducer was added if needed) and incubated at 37°C for 3 hours. Cells were then harvested by excising individual spots from the agar plates into LB medium. For liquid (well-mixed) conditions, mixtures of attacker and prey cells were established then incubated at 37°C, 180rpm for 2, 4, 6, 8, 24, 48 hrs. Suspensions were serially diluted and plated on selective media (selection for prey cells) for quantification of CFUs, then assessed the survival of prey cells. Survival rate was calculated as:

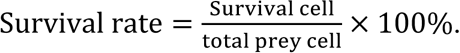

Killing efficiency was calculated as:

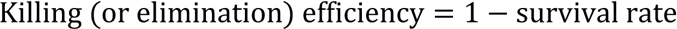

### SimCell purification and characterisation

For T6SS-assembled SimCells, overnight cultures were transferred into fresh LB medium with corresponding antibiotics at a ratio of 1:100 and grown at 37 °C, 180 rpm to an OD600 = 1, inducer was added when OD600 reached 0.6 if needed. For NahG-expressing SimCells, overnight cultures were directly used. Cells were collected by 1500 g centrifuge for 10 min at room temperature, washed twice, and resuspended with M9 + 0.2% casamino acids media, inducer was added if needed. 1μM of crystal violet was added to induce SimCell conversion. Incubated overnight at 37 °C, 180 rpm. Then ceftriaxone (100 μg/mL), penicillin G (100 μg/ mL), and cefotaxime (100 μg/mL) were added to the culture and further incubated at 37 °C, 180 rpm for 4 hours. 100 μL of cultures were plated for purity check and the rest were kept at 4 °C for further use. Cultures were washed twice with 1×PBS to remove the crystal violet and antibiotics before being used for bacterial killing assay.

### Growth arrest assay

Overnight cultures were diluted in 1:1000 ratio in 200 μL LB with corresponding antibiotics in a flat 96-well plate. The plate was sealed with a Breathe-Easy sealing membrane and incubated with Synergy 2 microplate reader (BioTek) at 37 °C, 1000 rpm. OD600 was measured every 15min since incubation. After 2-3 hours, OD600 reached ∼0.3, 0.2 μL of crystal violet to induce the SimCell conversion. OD600 were continuously measured for over 35 hours.

### Plating assay

The culture (5 μL) from each group was extracted and dot-plated in LB agar without antibiotics. Plates were incubated for 24 hr at 37 °C then images were collected to record the colony growth conditions.

### Mini-SimCell production and purification

For T6SS-assembled mini-SimCells, overnight cultures were 1:100 inoculated into 100mL fresh LB medium with corresponding antibiotics and grown at 37 °C, 180 rpm to an OD600 = 1, inducer was added when OD600 reached 0.6 if needed. For NahG-expressing SimCells, 100mL of overnight cultures were directly used. The culture was centrifuged at 2000 g for 10 min at 4 °C to remove the normal-size cells, the supernatant was retained for further centrifugation at 12,000 g for 15 min at 4 °C. The pellet was resuspended in 1 mL of fresh LB and pooled together. Ceftriaxone (100 μg/mL), penicillin G (100 μg/ mL), and cefotaxime (100 μg/mL) were added to the culture and incubated at 37 °C, 180 rpm overnight. The culture was first centrifuged at 500 g for 10 min at 4 °C to remove cell debris. Then the supernatant was retained and further centrifuged at 12,000 g for 15 min at 4 °C to collect the mini-SimCells. The final pellet was resuspended in 1 mL of 1xPBS solution and stored at 4 °C until further use. Cultures were washed twice with 1xPBS to remove the antibiotics before being used for bacterial killing assay. 5uL of each sample was used for plating assay to verify the purity.

### Mini-SimCell size distribution and concentration characterisation

The size distribution and concentration of Mini-SimCells were characterised by nanoparticle tracking analysis (NTA) using ZetaView (Particle Metrix, Inning am Ammersee, Germany). Prior to the measurement, Silica 100 nm microspheres (Polysciences Inc., Philadelphia, United States) were used for the quality check. Purified mini-SimCells were diluted in 1xPBS to an optimal concentration. The same batch of 1xPBS solution was used as the control group. 1mL of each sample was loaded onto the sample chamber using a syringe to avoid introducing bubbles. Background 1×PBS was used for flushing away the previous sample before loading a new sample. In the sample chamber, mini-SimCells were illuminated by a laser beam, the scattered light was captured, and the movement of each particle over time was tracked by a live camera. Based on Brownian motion, the diffusion coefficient of the particles was calculated, and then the size of each particle was determined using the Stokes-Einstein equation. Mini-SimCell concentrations were calculated by the number of particles in a known volume compared to the control group. Each sample was measured from 11 positions in 2 cycles. Parameter set as: sensitivity = 80, frame = 30, shutter speed = 100. For T6SS Mini-SimCells, a 500 nm filter was added to observe the fluorescent particles selectively.

### Multi-dose dual-mechanism mini-SimCell consortium Killing Assay

*E. coli* with and without surface-displayed EPEA epitope were designated as Ag+ and Ag- strains respectively. Both strains were grown overnight in LB medium at 37 °C with shaking at 180 rpm, then normalised to an OD600=1.25. Equal volume (20 µL) of each culture was co-inoculated into 1 mL of fresh LB in 1.5 mL Eppendorf tubes for 24 hrs to establish a steady status of the mixed population. Following incubation, 20 µL of each co-culture was collected for serial dilution and plated on selective plates for prey cells and counted colony forming units (CFUs), defining this as Day 0. Aspirin was then added into LB medium with a final concentration of 800 µM. Fresh mini-SimCells were prepared daily for every treatment dose. The engineered dual-mechanism ++ mini-SimCells consisted of Attaker1: mini-SimCell Nb-T6SS and Attaker2: mini-SimCell Nb-NahG at an n:n ratio of 1:1. The non-engineered - - mini-SimCells were served as negative control. For each dose, 100 µL 10^12^ ++ / - - mini-SimCells were well-mixed and then added into the co-culture. The first dose of ++ / - - mini-SimCells was added after initial 24-hr incubation, followed by an additional 16 hrs incubation at 37 °C, 180 rpm. Afterwards, 20 µL of the co-cultures were collected for plating and CFU counting (Day 1). The co-cultures were then diluted 1:50 into fresh LB medium containing 800 µM aspirin, and incubated at 37 °C, 180rpm for 8 hrs. This process was repeated for the 2^nd^, 3^rd^, 4^th^ doses of mini-SimCells (Day 2,3,4).

### Double-color Flow Cytometry Assay for Binding Population Quantification

Overnight *E. coli* ST131 cultures was 1:100 inoculated in 5 mL M9 minimal media with 0.4% glucose and 1 mM EDTA, then grew at 37°C, 180 rpm for 48 hrs. BL21:Ag and BL21: Nb39-mRFP strains were 1:100 inoculated at 37°C, 180 rpm in 5 mL LB medium and grew overnight. All cultures were centrifuged at 1500 g for 10 min, washed twice and resuspended with 1mL 1xPBS solution. *E. coli* ST131 and BL21:Ag cultures were stained with 1uM of SYTO 9 dye at room temperature for 2 hrs in dark. Then all cultures washed twice with 1xPBS solution and resuspended with 1xPBS solution, OD600 normalised to 1.0. *E. coli* ST131/ BL21:Ag and BL21: Nb39-mRFP cultures were mixed in v:v / 1:1 ratio in total volume of 1.5mL. Mixed cultures were settled still in dark overnight. Each culture was 1:100 diluted with 1×PBS solution and used for FACS analysis. BD Aria Fusion was used for flow cytometry analysis.

### Data Availability Statement

The data generated in this study are provided in the main text and/or Supplementary information, where the source data are listed in the Source Data file. Source data are provided with this paper.

## Acknowledgements

W.E.H. thanks EPSRC (EP/M002403/1 and EP/Y014073/1) for financial support. Y.D. gratefully acknowledges support from Oxford Interdisciplinary Bioscience DTP, funded by BBSRC (BB/T008784/1). K.R.F. is funded by Wellcome Trust Investigator award 209397/Z/17/Z and European Research Council Grant 787932. We thank Harris Saeed from the Engineering Department, University of Oxford, for helping with the SEM. Sample process and image for SEM were conducted at the Sir William Dunn School of Pathology, University of Oxford. We would like to acknowledge Dr. Naveed Akbar from the Radcliffe Department of Medicine, University of Oxford, for providing ZetaView equipment and technical expertise, and Vasiliki Tsioligka from the Flow Cytometry Facility, Sir William Dunn School of Pathology, University of Oxford, for providing flow cytometry technical expertise. We also thank all members of the Huang lab for the discussions and support.

## Author Contributions Statement

W.E.H conceived the original idea; Y.D. and W.E.H. designed research; Y.D. and X.J. performed research; X.J. T.D. Y.W. and E.B. contributed new reagents/analytic tools. Y.D., K.R.F. and W.E.H. analysed and interpreted data; Y.D. and W.E.H. drafted the manuscript; W.E.H. and K.R.F. provided supervision; all authors reviewed and revised the manuscript.

