## Supplementary Information for "Reprogrammed SimCells for Antimicrobial Therapy"

Classification: Biological Sciences, Microbiology

Keywords: synthetic biology, SimCells, mini-SimCells, nanobody-antigen interaction, antibacterial therapy, type VI secretion system, aspirin, catechol, hydrogen peroxide, antimicrobial resistance

### Supplementary Results

#### ***Heterologous Expression, Assembly and Function of T6SS in E. coli***

The expression, assembly and function of T6SS were verified in *E. coli* BL21(DE3) parental cells. Following the confirmation of heterologous T6SS expression with fluorescence analysis, a bacterial killing assay was conducted to assess the functional efficacy of the T6SS in contact-promoting conditions (as in solid growth conditions). After a 4-hour co-incubation on the LB plate, both constitutive and induced T6SS-expressing attacker cells demonstrated significant killing activity against prey cells compared to the inactive controls. Notably, the induced T6SS expression resulted in the elimination of over 98% of the prey cells in the mixture (Figure S2a). These findings confirm the functional efficacy of heterologously expressed T6SS.

To examine the stability of T6SS machinery, flow cytometry analysis was conducted to monitor the fluorescence signals of sfGFP-labelled T6SS VipA sheath in constitutive and induced T6SS-expressing strains over a 96-hour time course. For the constitutive T6SS-expressing strain, over 97% of cells remained $GFP^+$ after 96 hours of continuous culture, and no significant decrease was observed in the duration. Notably, the proportion of $GFP^+$ cells in the induced T6SS-expressing strain increased progressively over time with the presence of the inducer in the culturing medium. However, there were fewer $GFP^+$ cells in the induced T6SS-expressing strain (~65%) compared to the constitutive group (Figure S2b). To further assess machinery functionality, we evaluated the competitive capacity of both attacker strains against a prey strain (*E. coli* DH10B) after 96 hours of growth. Significant out-competition of the prey cells was observed in both groups, with the constitutive T6SS-expressing strain exhibiting a higher inhibition efficiency (~46% elimination of prey cells) compared to the induced group (~37% elimination of prey cells) (Figure S2c). The results suggest a higher stability for T6SS under constitutive control, possibly due to a relatively lower burden for cell growth. A decline of the T6SS functionality was noticed after 96-hour culturing without a loss in the $GFP^+$ cell population, which may be attributed to the decrease in cell viability due to an insufficient energy supply.

#### ***Hydrogen Peroxide ($H_2O_2$) Produced from Catechol Polymerisation Shows Broad-spectrum Antimicrobial Activities***

Catechol is a broad-spectrum antimicrobial candidate (1–3). Through an auto-polymerisation process, $H_2O_2$ is generated and subsequently disrupts a broad range of bacterial cellular processes (Figure 6a) (4–7). A hydrogen peroxide assay with 800 µM catechol in M9 minimal medium showed sustained $H_2O_2$ release over 21 days, reaching concentrations above 300 μM (Figure S6a). In the disc diffusion assay, the formation of inhibition zones verified the antibacterial activity of 5 mM to 100 mM catechol against *Pseudomonas putida* (Figure S6b). Additionally, when different concentrations of catechol were added to *E. coli* DH10B cultures, significant inhibition was observed at 5 mM and higher concentrations, with a time-dependent increase in killing efficiency for the 5 mM group (Figure S6c). This indicates a prolonged release of $H_2O_2$ from catechol polymerisation, contributing to sustained antibacterial activity over time.

***Catechol is Generated and Released from Parental Cells and SimCells***

To achieve a sustained cellular conversion of aspirin to catechol, the NahG enzyme was integrated into our engineered parental cell, SimCell and mini-SimCell platforms, regulated by a strong constitutive promoter J23100 followed by a strong ribosome binding site (RBS) BBa_B0031. The NahG enzyme and mRFP labelling were positioned downstream of the nanobody (Figure S7a). As for the negative control (NC) strain, SalR, a regulatory protein for a salicylate hydroxylase SalA, which shares high function similarity with NahG, is used as substitution of NahG.

With 800 μM aspirin added into the NahG+ parental cells, the filtered supernatant from the overnight culture showed a dark-brown colour (Figure S7b), indicating the cell-produced catechol was permeable to the cell membrane and was released to the surrounding environment. To test the antimicrobial effect, the collected supernatants were subsequently mixed in varying ratios with concentrated LB medium to normalise the nutritional content. These supernatant-LB mixtures were then utilised as the growth medium for the prey cells (*E. coli* DH10B). As shown in Figure S7b, the medium containing 20% NahG+ supernatant demonstrated a significant inhibitory effect compared to the control group. The cell growth was reduced in mixtures with 50% and 80% supernatant, likely due to the accumulation of secondary metabolites from prior culturing. Similar results were observed using supernatant from NahG+ SimCells (Figure S7c). This result corroborates that catechol is permeable across the cell membrane to exert its antimicrobial activity.

### Supplementary Methods

#### Assessing T6SS machinery stability with flow cytometry

Flow cytometry was applied to track the T6SS-related fluorescence in a single cell. Overnight cultures were 1:100 re-inoculated into the fresh LB medium with antibiotics for selection. All cultures were continuously incubated at 37°C, 180 rpm. Samples were collected at 24, 48, 72, 96h, centrifuged and resuspended in 1×PBS. A FACS Calibur (BD Biosciences) with an FL1 filter (FITC channel) was used for flow cytometry analysis. CellQuest was used to acquire 100,000 viable cells' fluorescence data for each experiment. Data analysis was performed by FlowJo software.

#### SYTO 9 staining assay

200 µL of culture from each group were collected, washed twice then resuspended with 1×PBS solution. A final concentration of 100nM of SYTO 9 stain was added and the cultures were incubated in a dark place at room temperature for 60 min. Cell pellets were collected and washed with 1 mL sterile $ddH_2O$ three times to remove the dye. 2 µL of the sample was used for fluorescence microscopy. Excitation/emission for SYTO 9 stain was 485/498 nm.

#### Single-cell Raman Spectra (SCRS)

Raman microspectroscopy was used for the characterisation of single bacterial cells' metabolic profiles. Prior to the measurement, the SimCells were washed with sterile $ddH_2O$ three times to remove the culture medium and extracellular metabolites and then diluted until individual cells could be observed under the microscope. 2 µL of each sample was dropped and air-dried onto an aluminium-coated slide. Raman spectroscopic acquisition was performed using a LabRAM HR Evolution confocal Raman microscope with a 100×/0.75 air objective (HORIBA, UK). A 532-nm neodymium-yttrium aluminium garnet laser with a 300-grooves $mm^{-1}$ diffraction grating was used for SCRS. ~80 mW laser power was set, and before focusing, it was attenuated by neutral density (ND) filters to protect the samples. 100–3200 $cm^{-1}$ spectrum was obtained and recorded by LABSPEC 6 software (HORIBA, UK). All raw spectra were corrected using cosmic ray correction, pre-processed using polyline baseline fitting and subtraction, and vector-normalised for the entire spectrum. For optimising single-cell level visualisation, Linear discriminant analysis (LDA) was applied for SCRS dimension reduction. R studio was used for data analysis and figure plotting. Raman peak at 785 $cm^{-1}$ (cytosine ring breathing mode) indicated the DNA/RNA level of the single cell.

**Fluorescence Microscopy**

Prepared fresh parental cell or purified SimCell / mini-SimCell samples were washed twice and then resuspended with sterile deionised water to remove the remaining medium. 2 µL of the sample was dropped and air-dried on the aluminium-coated slide. Nikon Ti Eclipse was used for fluorescence microscopy. Fiji was used for image analysis.

**Scanning Electron Microscopy (SEM)**

To prepare sterile coverslips, clean coverslips were dipped in 70% ethanol overnight, air-dried then exposed to UV light for 30 min. For coating, clean and sterile coverslips were dipped into 0.01% poly-L-lysine solution for 1-2 hours at room temperature, then washed twice with sterile water. The coated coverslips were allowed to dry in a 60°C drying hood overnight before use and were stored at 4°C in a sterile container for further use. For sample preparation, purified mini-SimCell culture was diluted into an appropriate concentration to avoid overlap between cells. The coated coverslips were dipped in the culture and incubated at 37°C for 30 minutes. For primary fixation, the coverslips were dipped into the 2.5% glutaraldehyde solution and fixed at room temperature for 1 hour. After primary fixation, the samples can be stored at 4°C for up to 1 week or proceeded to secondary fixation. For secondary fixation, the fixed samples were first rinsed with pH=7.2 0. 1M PIPES buffer for three times, 5 min each time. A sufficient volume of 1% $OsO_4$ in 0.1 M PIPES buffer was used to cover the samples. The samples were incubated at 4 °C for 1 hour and then rinsed with dd$H_2O$ three times, 5 min each time. Subsequent to secondary fixation, the samples were dehydrated using graded EtOH concentrations of 50%, 70%, 90%, and 95%, one time for 5 min each and finally three times using 100% EtOH for 10 min. For the drying step, the EtOH was removed and 0.5 mL hexamethyldisilazane (HMDS) was quickly added to each sample. After 3 min incubation, the HMDS was removed, and the samples were left for air dry overnight or at least 3 hours to allow the HMDS fumes to evaporate. This step was performed in the fume hood. Once dried, the samples were attached to carbon adhesive tape on an SEM stub, sputter-coated with gold/palladium, and then viewed using Zeiss Sigma 300 SEM. Fiji was used for image analysis.

**$H_2O_2$ generation assay**

The concentration of hydrogen peroxide is measured using a fluorimetric hydrogen peroxide assay kit (Sigma-Aldrich MAK165). 10mL of 800 µM catechol in M9 minimal medium and blank M9 minimal medium is prepared and continuously incubated at 37 °C, 180rpm. At 1, 3, 7, 11 and 21 day, 50 µL of each sample is collected and mixed with 50 µL of the Master Mix in each well of 96-well plate (Greiner, black). All samples were run in triplicates. The plate was incubated at room temperature for 30 min in the dark. The fluorescence intensity is measured at λex = 540/ λem = 590 nm using a fluorescence plate reader (Tecan Spark). Gain = 60. All readings were corrected by subtracting the background. The hydrogen peroxide concentration for the samples is calculated based on the standard curve.

**Disc diffusion method for antimicrobial activity characterisation of catechol**

This method is adapted from Kirby-Bauer disc diffusion method for antibiotic susceptibility testing. An overnight bacterial suspension was prepared and normalised to $OD_{600}$ =2.0. Sterile L-shaped spreaders were used to evenly spread the bacterial suspension over the surface of fresh LB agar plates. Allowing the inoculum to dry for a short period, small paper discs impregnated with 50 µL of 0/ 5 mM/ 10 mM/ 0.1 M catechol were placed on the surface of the agar plates. Sterile forceps were used for placing the discs. Triplicates were applied for each plate with even spacing between discs. All agar plates were incubated at 37°C for 24 hours before capturing the images. Inhibition zones appeared as clear, circular areas around each disc. Fiji was used for image analysis and measurement of inhibition zones.

**Assessing permeability and extracellular antimicrobial activity of cell-produced catechol**

Overnight cultures were 1:100 re-inoculated into fresh LB medium with antibiotics for selection, grew until stationary phase and centrifuged to collect all cells. Cell pellets were washed twice with 1×PBS to remove traces of culture medium and antibiotics, then resuspended with fresh minimal medium (M9 medium) with 800uM Aspirin. All cultures were normalised to a volume of 50 mL and $OD_{600}$ of 2.0. After overnight incubation at 37°C, 180 rpm, the supernatants were collected, filtered for sterilisation, and mixed in different ratios with concentrated fresh LB medium to normalise the nutrition level. The final supernatant-LB mediums contained 20%, 50%, 80% of the supernatant respectively and 1xLB. Then these media were used as the growth medium for prey cells (*E coli* DH10B). Overnight cultures of prey cells were 1:1000 re-inoculated into 200 µL of different growth media with corresponding antibiotics in a flat 96-well plate. The plate was sealed with Breathe-Easy sealing membrane and incubated with a Synergy 2 microplate reader (BioTek) at 37 °C, 1000 rpm. $OD_{600}$ was measured every 15 min for 18 hours.

### Supplementary Tables

**Table S1.** Comparison of advantages and challenges between various antimicrobials.

| | Approach | Biosafety | Stability | Resistance risk | Manufacture | Refs |
|---|---|---|---|---|---|---|
| Traditional antimicrobials | Antibiotics | √ | √ | High | √ | (8–10) |
| Non-traditional antimicrobials | Antimicrobial peptides (AMPs) | 70% of AMPs are predicted to have hemolytic activity | Instable, with short half-lives | Medium | √ | (11–13) |
| | Bacteriophage | √ | √ | High | Difficult | (14–17) |
| | Monoclonal antibodies | √ | √ | Medium | Complex and Expensive | (18–20) |
| | Nanoparticles (NPs) | Potential toxic effects in human health and environment | Appropriate stabilizer required | Low | Can be difficult | (21–23) |
| | Gas therapy | Can be toxic | Uncontrollable, reliable delivery system to-be-developed | Low | √ | (24–26) |
| | Live biotherapeutic products (LBPs) | Biosafety and biocontainment concerns | √ | Medium to low | √ | (27–30) |

**Table S2.** Strains and plasmids used in this study.

| Strain | Description | Source |
|---|---|---|
| *Escherichia coli* DH5a | | Lab collection |
| *E. coli* DH10B | | Lab collection |
| *E. coli* BL21(DE3) | | Lab collection |
| *E. coli* BL21(DE3) ΔminD | | Lab collection |
| *Pseudomonas putida* UWC1 | | Lab collection |
| *E. coli* ST131 | | NCTC 13441 |
| BL21: Ag | *E. coli* BL21(DE3): pDSG287_sfGFP | This study |
| BL21: Nb | *E. coli* BL21(DE3): pDSG289 + pL1_WH002_EF | This study |
| BL21: Null | *E. coli* BL21(DE3): pDSG291 + pL1_WH002_EF | This study |
| BL21: constitutive T6SS | *E. coli* BL21(DE3): pT6S_NP | This study |
| BL21: induced T6SS | *E. coli* BL21(DE3): pT6S_Tet | This study |
| BL21: Nb+T6SS+ | *E. coli* BL21(DE3): pDSG289 + pT6S_NP | This study |
| BL21: Nb+T6SS- | *E. coli* BL21(DE3): pDSG289 + pT6S_N3-NP | This study |
| BL21: Null T6SS+ | *E. coli* BL21(DE3): pDSG291 + pT6S_NP | This study |
| BL21: Null T6SS- | *E. coli* BL21(DE3): pDSG291 + pT6S_N3-NP | This study |
| BL21: Nb+NahG | *E. coli* BL21(DE3): pNb_NahG | This study |
| BL21: Nb39_mRFP | *E. coli* BL21(DE3): pNb39_NahG | This study |
| SimCell | *E. coli* BL21(DE3): pRH12x | This study |
| Nb-SimCell | *E. coli* BL21(DE3): pRH12x + pDSG289 | This study |
| Null-SimCell | *E. coli* BL21(DE3): pRH12x + pDSG291 | This study |
| SimCell: constitutive T6SS | *E. coli* BL21(DE3): pRH12x + pT6S_NP | This study |
| SimCell: induced T6SS | *E. coli* BL21(DE3): pRH12x + pT6S_Tet | This study |
| SimCell: Nb+T6SS+ | *E. coli* BL21(DE3): pRH12x + pDSG289 + pT6S_NP | This study |
| SimCell: Nb+T6SS- | *E. coli* BL21(DE3): pRH12x + pDSG289 + pT6S_N3-NP | This study |
| SimCell: Null T6SS+ | *E. coli* BL21(DE3): pRH12x + pDSG291 + pT6S_NP | This study |
| SimCell: Null T6SS- | *E. coli* BL21(DE3): pRH12x + pDSG291 + pT6S_N3-NP | This study |
| SimCell: Nb+NahG | *E. coli* BL21(DE3): pRH12x + pNb_NahG | This study |
| Mini-SimCell (Mini-SimCell Null) | *E. coli* BL21(DE3) ΔminD | This study |
| Mini-SimCell: constitutive T6SS | *E. coli* BL21(DE3) ΔminD: pT6S_NP | This study |
| Mini-SimCell: induced T6SS | *E. coli* BL21(DE3) ΔminD: pT6S_Tet | This study |
| Mini-SimCell: Nb+T6SS+ | *E. coli* BL21(DE3) ΔminD: pDSG289 + pT6S_NP | This study |
| Mini-SimCell: Nb+T6SS- | *E. coli* BL21(DE3) ΔminD: pDSG289 + pT6S_N3-NP | This study |
| Mini-SimCell: Null T6SS+ | *E. coli* BL21(DE3) ΔminD: pDSG291 + pT6S_NP | This study |
| Mini-SimCell: Null T6SS- | *E. coli* BL21(DE3) ΔminD: pDSG291 + pT6S_N3-NP | This study |
| Mini-SimCell: Nb+NahG | *E. coli* BL21(DE3) ΔminD: pNb_NahG | This study |

| | | |
|---|---|---|
| Mini-SimCell Nb39-T6SS | *E. coli* BL21(DE3) ΔminD: pTet_Nb39 + pT6S_NP | This study |
| Mini-SimCell Nb39-nahG | *E. coli* BL21(DE3) ΔminD: pNb39_NahG | This study |

| **Plasmid** | **Description** | **Source** |
|---|---|---|
| pDSG287 | Plasmid with surface-displayed EPEA | Addgene #115606 (31) |
| pDSG287_sfGFP | Plasmid with surface-displayed EPEA and sfGFP | This study |
| pDSG289 | Plasmid with surface-displayed anti-EPEA | Addgene #115608 (31) |
| pDSG291 | Plasmid with surface-displayed Null | Addgene #115605 (31) |
| pL1_WH002_EF | Plasmid with constitutive high expression of mRFP | Lab collection |
| pT6S_NP | Plasmid with T6SS gene cluster from *Aeromonas dhakensis* under native constitutive promoter | Gift from Tao Dong, Southern University of Science and Technology, China (32) |
| pT6S_Tet | Plasmid with T6SS gene cluster from *Aeromonas dhakensis* under tetracycline-inducible promoter | (32) |
| pT6S_N3-NP | Plasmid with inactive T6SS gene cluster from *Aeromonas dhakensis* under native constitutive promoter | (32) |
| pT6S_N3-Tet | Plasmid with inactive T6SS gene cluster from *Aeromonas dhakensis* under tetracycline-inducible promoter | (32) |
| pRH12x | Plasmid with I-CeuI endonuclease | Lab collection(33) |
| pNb_NahG | Plasmid with constitutive high expression of anti-EPEA, salicylate hydroxylase NahG and mRFP | This study |
| pTet_Nb39 | Plasmid with surface-displayed Nb39 | This study |
| pNb39_NahG | Plasmid with constitutive high expression of Nb39, salicylate hydroxylase NahG and mRFP | This study |

**Table S3.** Primers used in this study.

| **Primer** | **Sequence 5' to 3'** |
|---|---|
| EPEA FWD | ATTTCACACAGGAACCTGACGTCTAAGAAACC |
| EPEA REV | CATCAAAAGATACATCAGAGCTTTTACGAG |
| EPEA-sfGFP FWD | CAGGAAAGAAACCATTATTATCATGACATTAAC |
| EPEA-sfGFP REV | CATCAAACATCAAGGGAAAACTGTCCATATGC |
| AmpR-mid-F | GATCCAGTTCGATGTAACCCACTCG |
| intimin-mid-R | CGGCAACAGTTTCACCAGTTTTCAAC |
| intiminN-test-F | GTCGAATGGTCAAGTTGTCGACCAG |
| Ag/Nb-seq-R | GCTGTTCACCATGAACAGATCGACAATG |
| nahG-Test-R | TCAACCAGATGTTCGTCGATCCC |
| mRFP-test-F | AGCTTCCACCGAACGTATG |

| | |
|---|---|
| vgrG-Seq-R | GTGCTCGTCATGCTTGATAT |
| vipA-Seq-F | GCCCAAAGAACGTATCAATATC |
| SP9 | ATGTCAAACTTTATACTTAAACCGG |
| SP10 | TGAGCTGCTTCTAAATTAGGAAA |
| SPGG_2 FWD | CAGATAAAAAAAATCCTTAGCTTTCGCT |
| SPGG_3 REV | TAACCGTATTACCGCCTTTGAGTG |

### Supplementary Figures

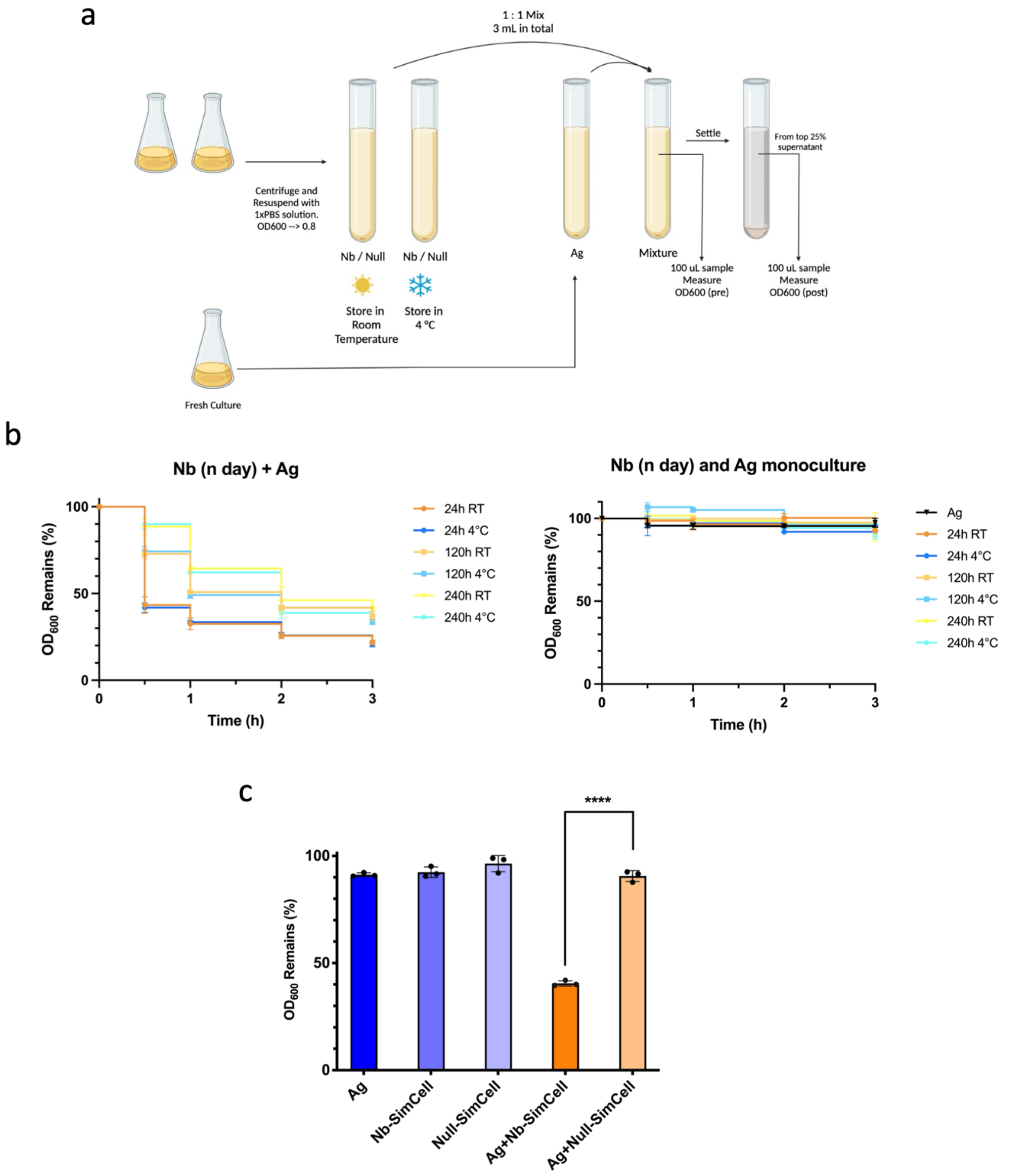


**Figure S1. Stability of surface-displayed nanobody. (a)** Macroscopic aggregation assay allows quantitative measurement of binding strength via optical density ($OD_{600}$). Adapted from previous work(31). **(b)** Macroscopic aggregation tests confirm that cells stored at room temperature (RT) or 4°C maintained the nanobody activity over 10 days. **(c)** SimCell Ag+Nb paired mixture shows significant aggregation-caused settling compared to unmixed and non-adhesion (Ag+Null) conditions after 3 hrs. For **(b)** and **(c)**, all cultures were normalised to the same initial $OD_{600}$ and mixed in 1:1 v/v ratio. N = 3 replicates, Error bars, ±1 SD, ****$p < 0.0001$ according to a 2-tailed paired t-test.

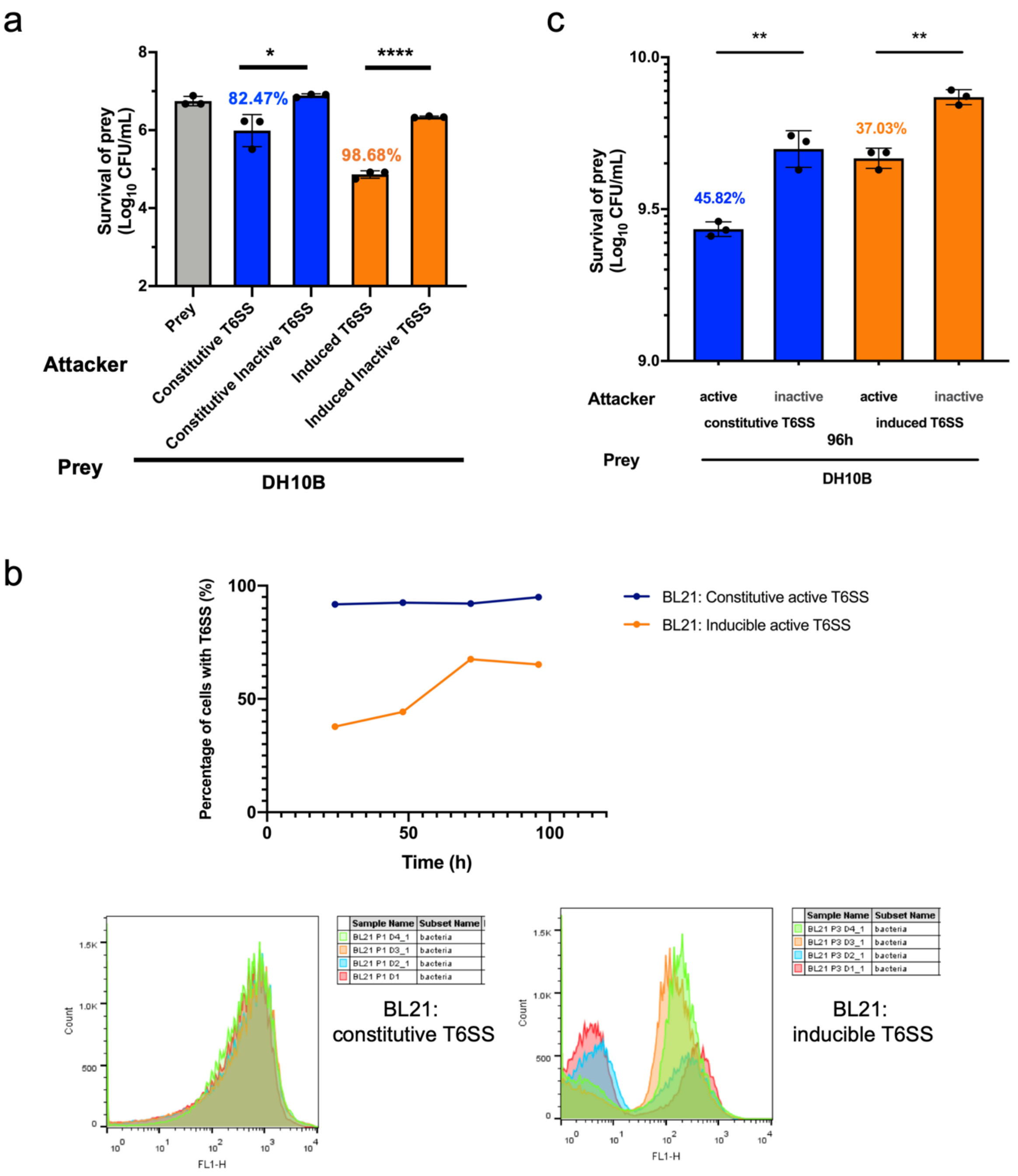


**Figure S2. Function and stability of heterologous T6SS in *E. coli* BL21(DE3) cells. (a)** Significant killing in solid condition indicates both constitutive and induced T6SS activity. **(b)** Flow cytometry results showing the GFP-labelling cells (T6SS-expressed cells) among the population and the change after continuous incubation. >97% of cells with constitutive expressed T6SS are stable for at least 96h. **(c) Killing against prey cells in solid condition indicates** after 96h of continuous incubation, both constitutive and induced expressed-T6SS BL21 parental cells maintain certain activity. N = 3 replicates, Error bars, ±1 SD, $*p < 0.1$, $**p < 0.01$, $****p < 0.0001$ according to a 2-tailed paired t-test. Killing efficiency was calculated as 1-survival cells/ total prey cells.

a
b
c

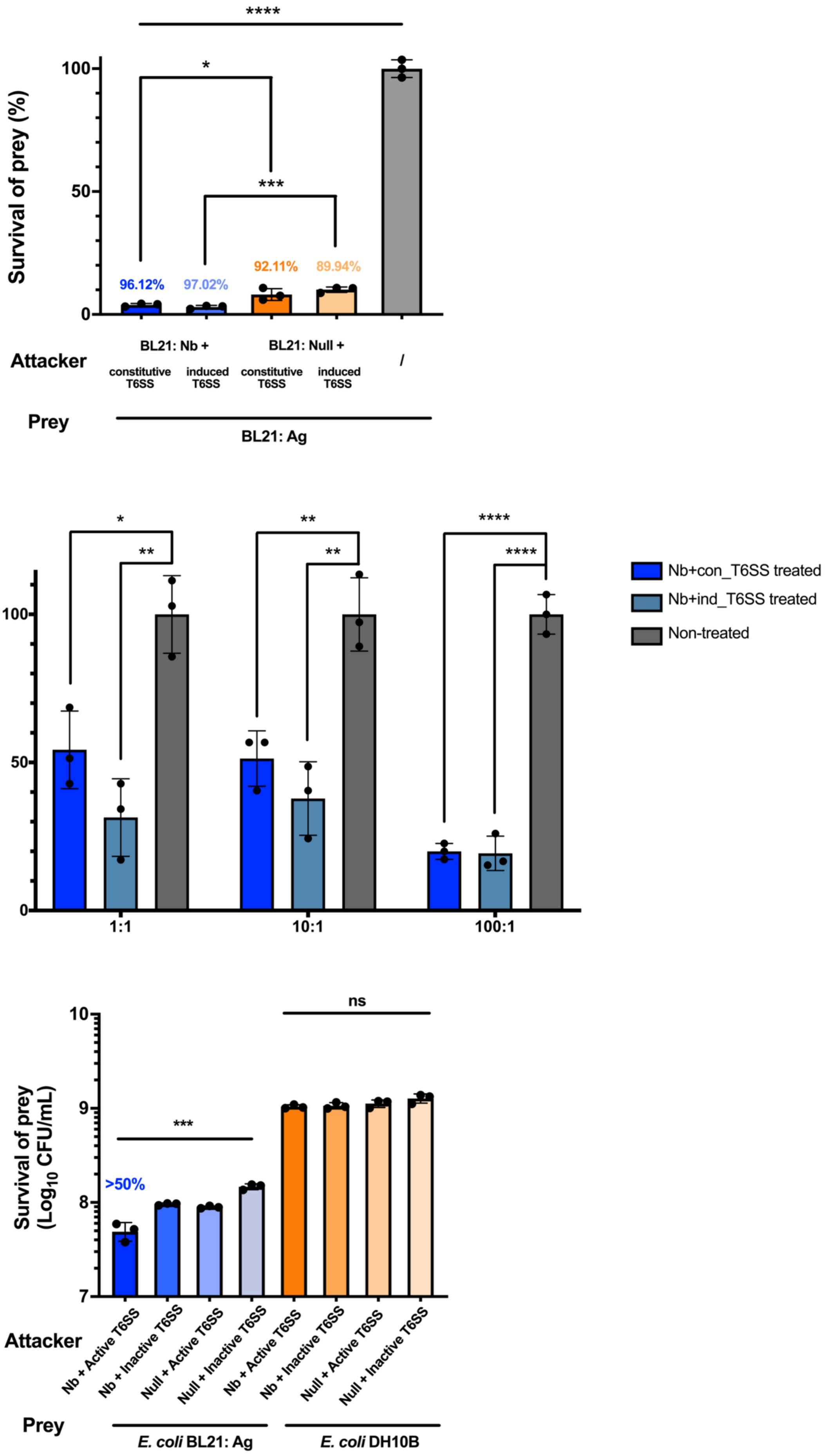
****
100
*
Survival of prey (%)
50
***
96.12%
97.02%
92.11%
89.94%
0
BL21: Nb +
BL21: Null +
Attacker
constitutive T6SS
induced T6SS
constitutive T6SS
induced T6SS
/
Prey
BL21: Ag
*
**
**
**
****
****
Nb+con_T6SS treated
Nb+ind_T6SS treated
Non-treated
Survival rate of prey (%)
100
50
0
1:1
10:1
100:1
10
ns
Survival of prey (Log10 CFU/mL)
9
***
>50%
8
7
Attacker
Nb + Active T6SS
Nb + Inactive T6SS
Null + Active T6SS
Null + Inactive T6SS
Nb + Active T6SS
Nb + Inactive T6SS
Null + Active T6SS
Null + Inactive T6SS
Prey
E. coli BL21: Ag
E. coli DH10B

207

**Figure S3. Nanobody-antigen binding enhanced T6SS in killing efficiency and enabled T6SS with killing specificity.** **(a)** Nanobody-antigen interaction further improves T6SS-mediated killing in solid condition. **(b)** Significant inhibition efficiency was observed with a lower to attacker: prey = 1:1 n:n ratio of Nb-T6SS BL21 parental cells as attacker cells in liquid condition. The efficiency was improved by increasing n:n ratio. **(c)** Nb-Ag recognition enables a specific T6SS-killing effect in liquid condition. For **(a)** to **(c)**, N = 3 replicates, Error bars, ±1 SD, *$p < 0.1$, **$p < 0.01$, ***$p < 0.001$, ****$p < 0.0001$ according to a 2-tailed paired t-test. Survival rate was calculated as survival cells/ total prey cells. Killing efficiency was calculated as 1-survival rate.

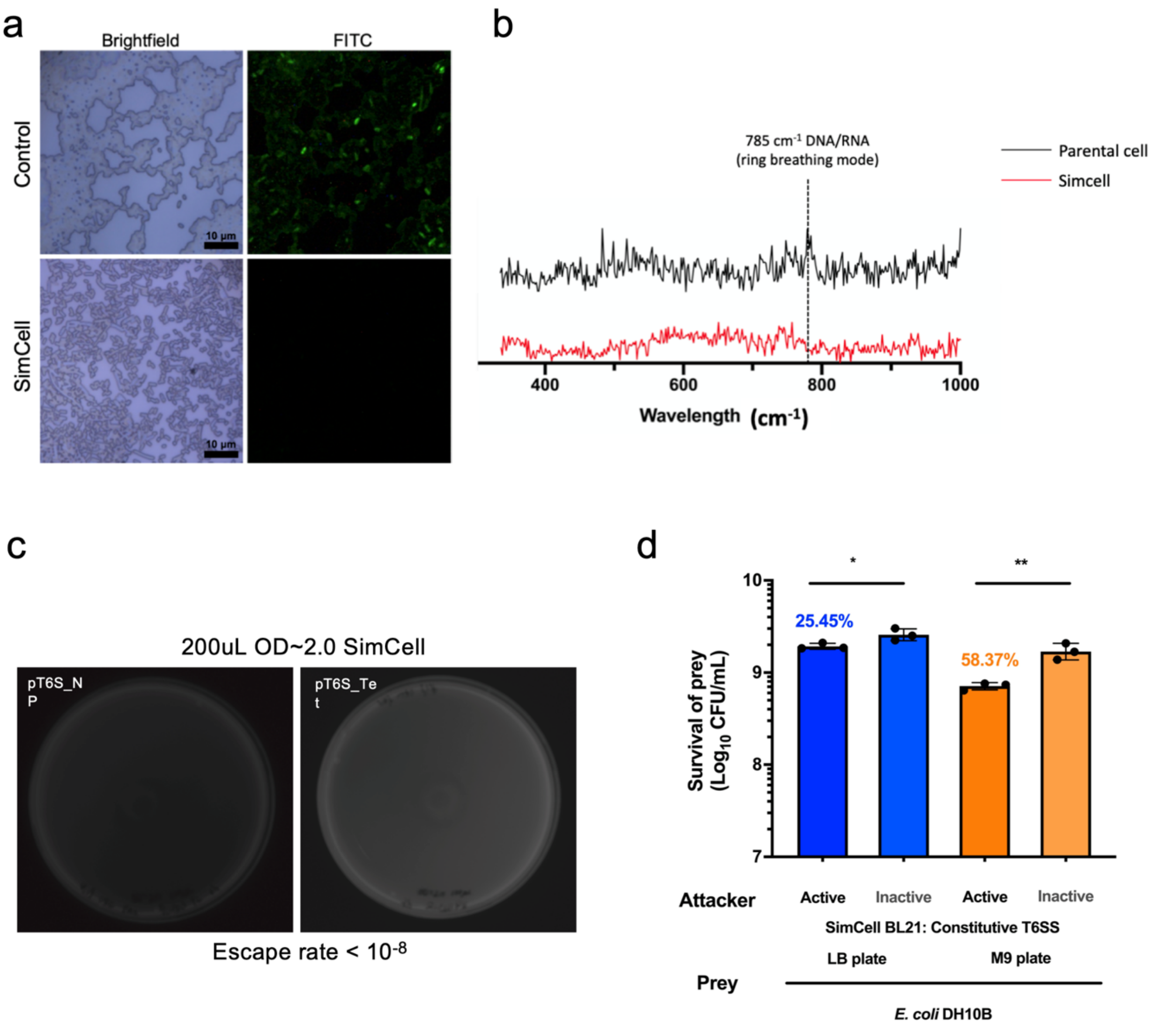


**Figure S4. Characteriasation of T6SS-assembled SimCells. (a)** SYTO 9 stain assay confirms successful SimCell conversion at single-cell level. Green fluorescence can be observed in control cells due to double-strand chromosomes being stained while negligible in SimCells. Scale bar, 10 µm. **(b)** Raman spectrum analysis confirms DNA reduction in SimCells. DNA/RNA peak in 785 $cm^{-1}$. **(c)** No colony is observed from plating 200 uL of $OD_{600}$ = 2.0 SimCell culture in LB agar plates, confirming the escape rate of lower than $10^{-8}$. All cultures were plated on LB agar plates and incubated for 24 hours at 37 °C. **(d)** Function tests of T6SS-assembled Simcells in solid condition. Incubation time = 3h. Attacker : prey v/v ratio = 10:1. N = 3 replicates, Error bars, ±1 SD, *p < 0.1, **p < 0.01 according to a 2-tailed paired t-test. Killing efficiency was calculated as 1-survival cells/ total prey cells.

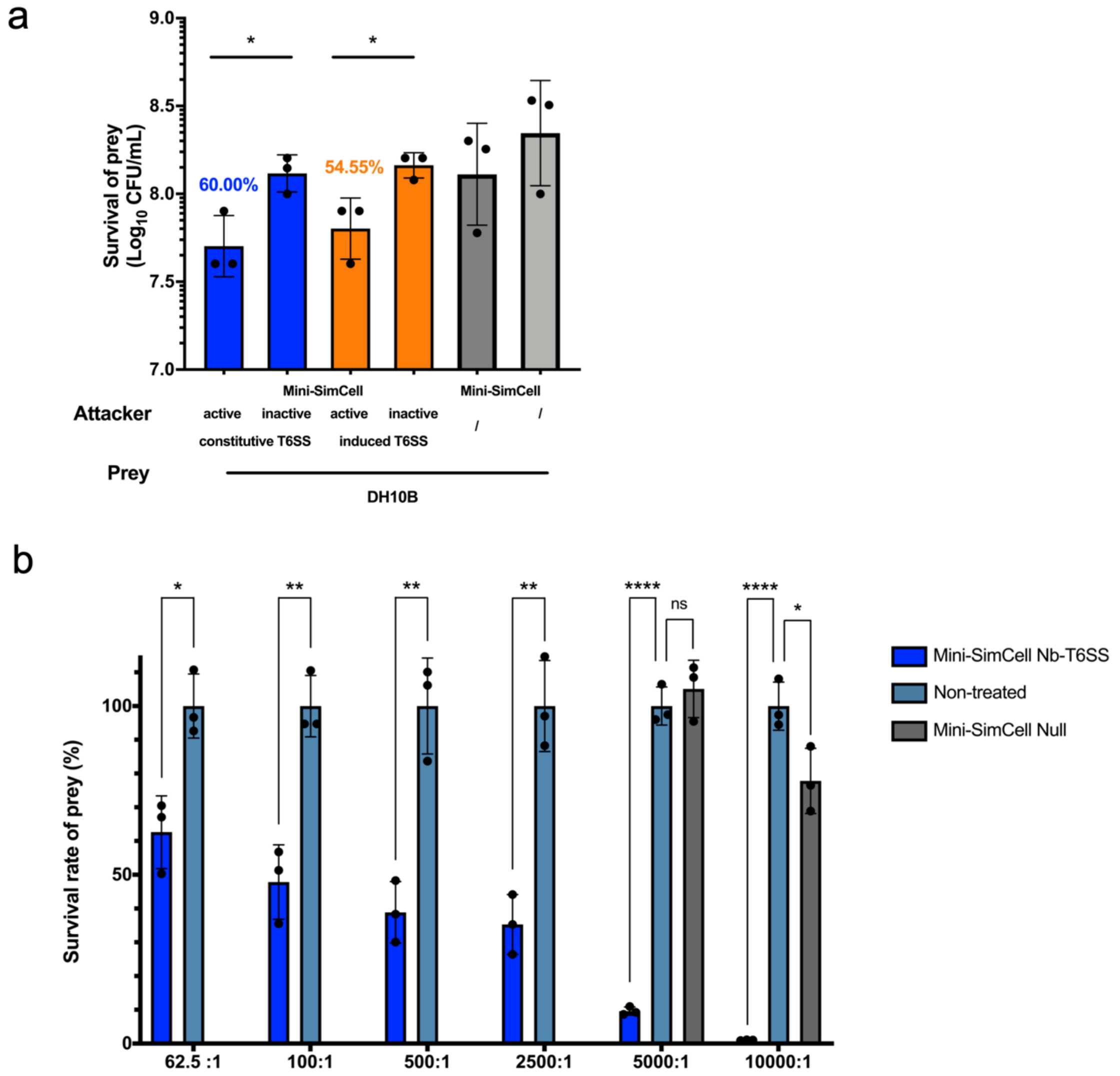


**Figure S5. Functional characterisation of T6SS-assembled mini-SimCell. (a)** Function tests of T6SS-assembled mini-Simcells in solid condition. Incubation time = 3h. Attacker : prey v/v ratio = 10:1. **(b)** Limit of killing (LOK) of Nb-T6SS mini-SimCells in liquid condition. Attackers (mini-SimCell Nb-T6SS) were mixed in liquid condition with prey cells in n:n ratio of Attacker : prey = 62.5/100/500/2,500/5,000/10,000:1, total Attaker cells were $1*10^{10}$, the maximum tolerated dose (MTD) reported in the previous human trial. Incubation time = 6h. Significant inhibition efficiency was observed in all groups. The efficiency was improved by increasing n:n ratio. For **(a)** and **(b)**, N = 3 replicates, Error bars, ±1 SD, $*p < 0.1$, $**p < 0.01$, $****p < 0.0001$ according to a 2-tailed paired t-test. Survival rate was calculated as survival cells/ total prey cells. Killing efficiency was calculated as 1-survival rate.

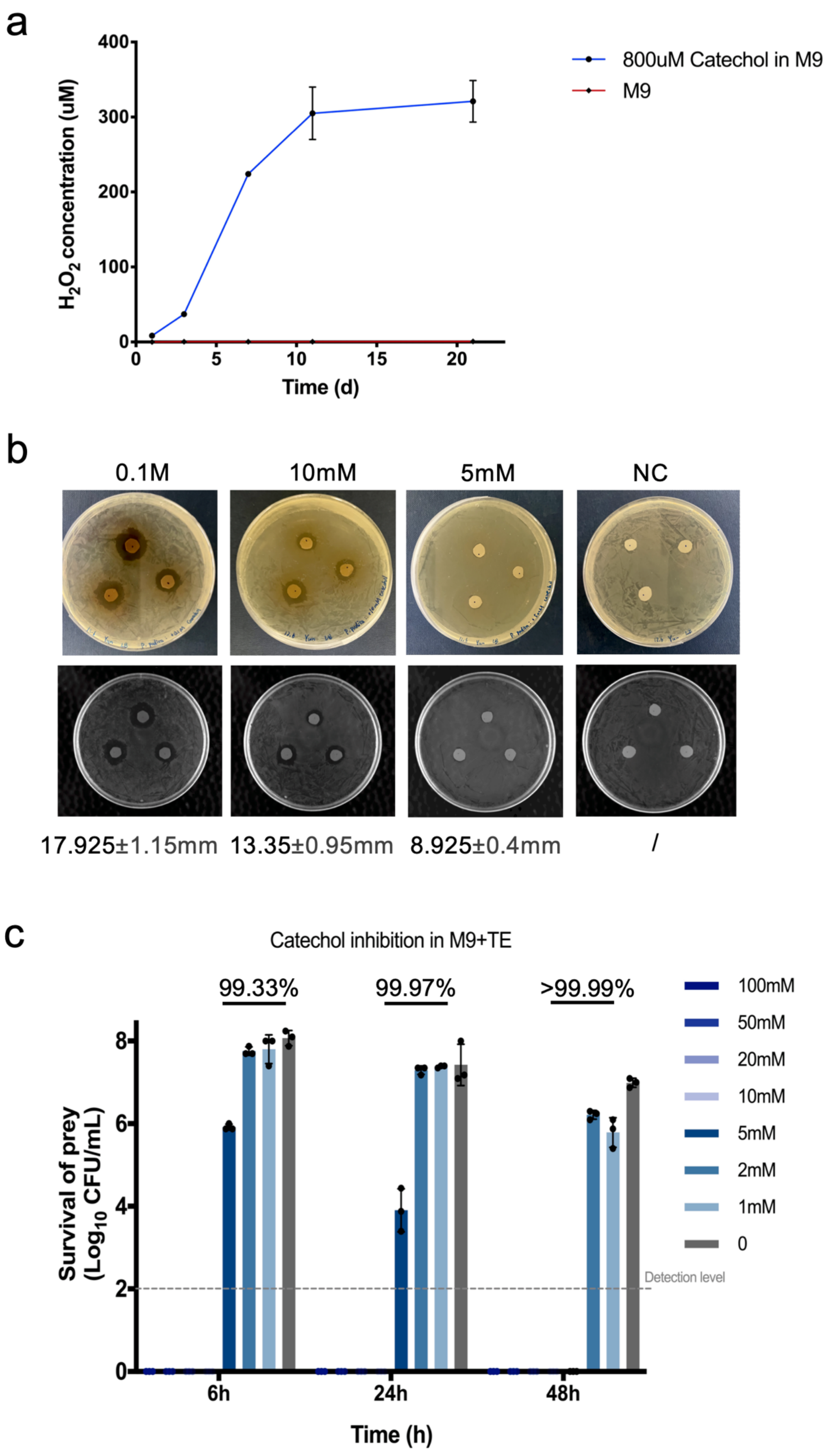
a
400
300
200
100
0
H2O2 concentration (uM)
0
5
10
15
20
Time (d)
800uM Catechol in M9
M9
b
0.1M
10mM
5mM
NC
17.925±1.15mm
13.35±0.95mm
8.925±0.4mm
/
c
Catechol inhibition in M9+TE
99.33%
99.97%
>99.99%
100mM
50mM
20mM
10mM
5mM
2mM
1mM
0
8
6
4
2
0
Survival of prey (Log10 CFU/mL)
Detection level
6h
24h
48h
Time (h)

**Figure S6. The antibacterial activity of catechol. (a)** $H_2O_2$ production over time from 800 μM catechol in 1×PBS at 37°C. Samples were taken at Day 1, 3, 7, 11 and 21. N = 3 replicates, Error bars, ±1 SD. **(b)** The antimicrobial activity of catechol was determined by the disc diffusion method. Values above the figures are the concentration of catechol. In negative control (NC) group, the equivalent volume of solvent for catechol solution preparation was used. Values below the figures are diameters of inhibition zones, DIZ (mm) ±1 SD, N = 3 replicates. **(c)** Inhibitory efficiency of different concentrations of catechol in liquid condition. 100/50/20/10/5/2/1mM of catechol was added into M9+TE (trace elements) medium and inoculated with $10^8$ prey cells (*E. coli* DH10B). After 6h, 24h and 48h incubation at 37°C, the survival of prey cells was determined by plating assay. N = 3 replicates, Error bars, ±1 SD. Killing efficiency was calculated as 1-survival cells/ total prey cells.

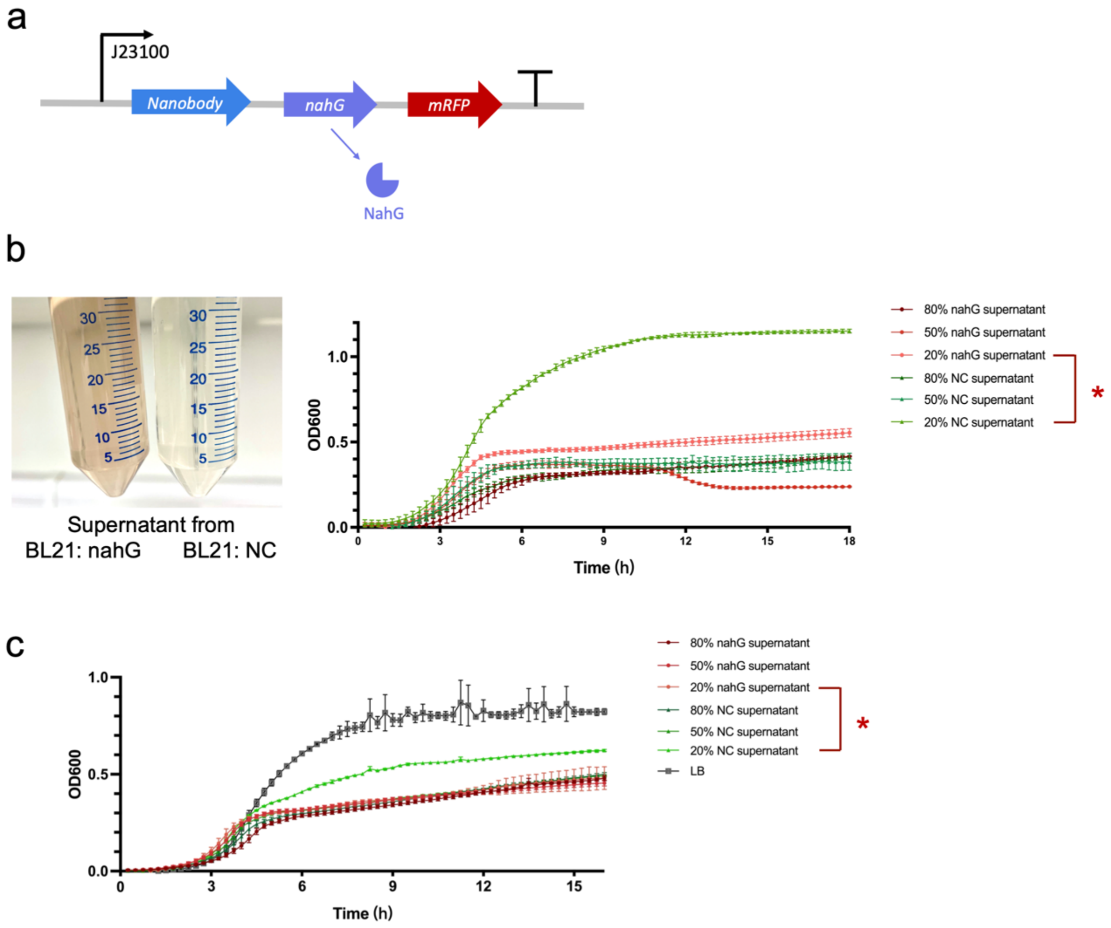


**Figure S7. The antimicrobial activity of cell-produced catechol. (a)** Engineered genetic circuit design for catechol generation. Nanobody and NahG enzyme are co-expressed under a strong constitutive promoter J23100. mRFP is co-expressed as an indicator of expression. In negative control (NC) group, SalR, a regulatory protein for a salicylate hydroxylase SalA, which shares high function similarity with NahG, is cloned in substitution of NahG. **(b)** Supernatant from 37°C overnight 800uM Aspirin + BL21: nahG/NC culture. The brown colour indicates the generation of catechol and its oxidation products. The supernatants were collected, filtered for sterilisation and mixed in different ratios with concentrated LB to normalise the nutrition level. Then the supernatant-LBs were used as the growth medium for prey cells (*E. coli* DH10B). 20% nahG supernatant showed a significant inhibition effect compared to the NC group. **(c)** Supernatant from 37°C overnight 800uM Aspirin + BL21-derived SimCell: nahG/NC culture. Similar results with the parental cells. For **(b)** to **(c)**, N = 3 replicates, Error bars, ±1 SD.

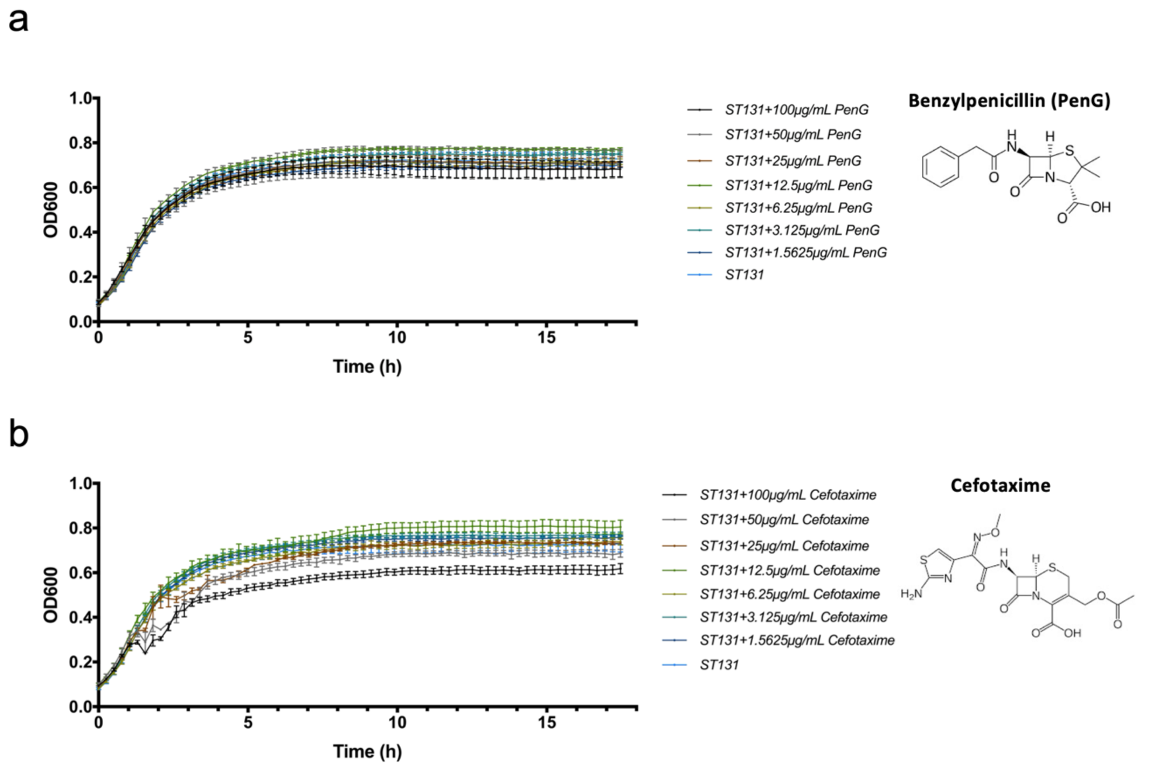


**Figure S8. *E. coli* ST131 strain exhibits multi-drug resistance. (a)** *E. coli* ST131 growth remains unaffected by benzylpenicillin (PenG) with concentration up to 100 μg/mL. **(b)** *E. coli* ST131 growth shows no significant inhibition by cefotaxime with concentration up to 100 μg/mL. N = 3 replicates, Error bars, ±1 SD.
